# Integrative Analysis of Spatial Transcriptome with Single-cell Transcriptome and Single-cell Epigenome in Mouse Lungs after Immunization

**DOI:** 10.1101/2021.09.17.460865

**Authors:** Zhongli Xu, Xinjun Wang, Li Fan, Fujing Wang, Jiebiao Wang, Wei Chen, Kong Chen

## Abstract

Immunological memory is key to productive adaptive immunity. An unbiased, high throughput gene expression profiling of tissue-resident memory T cells at their precise anatomical locations within the lung is fundamental to understanding lung immunity, but such spatial information has yet to be characterized. In this study, using a well-established *Klebsiella pneumoniae* infection model, we performed an integrative analysis of spatial transcriptome with single-cell RNA-seq and single-cell ATAC-seq on lung cells from mice after immunization using the 10x Genomics Chromium and Visium platform. We employed several deconvolution algorithms and established an optimized deconvolution pipeline to accurately decipher specific cell-type composition by anatomic location. We identified and located 12 major cell types by scRNA-seq and spatial transcriptomic analysis. Integrating scATAC-seq data from the same cells processed in parallel with scRNA-seq, we found epigenomic profiles provide more robust cell type identification, especially for lineage-specific T helper cells. When combining all three data modalities, we observed a dynamic change in the location of T helper cells as well as their corresponding chemokines for chemotaxis. Furthermore, cell-cell communication analysis of spatial transcriptome provided evidence of lineage-specific T helper cells receiving designated cytokine signaling. In summary, our first-in-class study demonstrated the power of multi-omics analysis to uncover intrinsic spatial- and cell-type-dependent molecular mechanisms of lung immunity. Our data provides a rich research resource of single cell multi-omics data as a reference for understanding spatial dynamics of lung immunization.

## INTRODUCTION

Immunological memory, consisting of B cell and T cell memory, is a key characteristic of adaptive immunity upon encountering with pathogen invasion. Typically, memory T cells can be classified into two categories: effector memory and central memory T cells. Effector memory T cells are able to produce effector cytokines and have cytotoxic activity against pathogen infected cells ^1^. Tissue-resident memory T cells (TRM) have more recently been defined as a new subset, predominantly residing in mucosal tissues, barrier surfaces, and other non-lymphoid organs but are also present in lymphoid sites ^2^. TRM are antigen experienced and are capable of rapidly responding to re-exposure to cognate antigen. TRM in the lung have been demonstrated to exhibit robust protective function against constant viral and bacterial challenge of the lungs and respiratory tract.

*Klebsiella pneumoniae* is an important cause of community-acquired pneumonia. In 2011, the U.S. National Institutes of Health Clinical Center experienced an outbreak of carbapenem-resistant *K. pneumoniae* that affected 18 patients, 11 of whom died. Thus, in addition to antimicrobial stewardship and hospital hygiene measures, there is a critical need for the development of novel therapeutic approaches to prevent and/or treat antibiotic resistant infections. Our previous work has demonstrated that *K. pneumoniae* specific Th17 cells are induced by immunization with whole bacterial lysate ^3^.These memory Th17 cells are both required and sufficient to provide serotype/antibody independent protection against a variety of strains of *K. pneumonia* including the recently described multidrug resistant New Delhi metallo-beta-lactamase strain.

Tissue localization of these TRM cells has been investigated thoroughly in the past and comparisons between mouse and human TRM have been characterized extensively using the well-established flow cytometric and transcriptomic approaches ^4^. However, an unbiased, high throughput gene expression profiling of TRM residing in various anatomical location within the lung, such as airway vs parenchyma, has not been possible using conventional flow cytometry, transcriptomic approaches, and even the most recently developed single-cell sequencing technology. These methodologies are limited as anatomic location specific information is lost after single-cell suspension is acquired.

Single-cell RNA-seq (scRNA-seq) has been gradually applied to study immune cells and immune responses in mouse lungs ^5–8^. Few recent studies utilize single-cell ATAC-seq (scATAC-seq) to measure the chromatin architecture of immune cells in mouse lungs ^9^. Although these technologies provide rich information in understanding cell heterogeneity and biology information of mouse lung, spatial information of single cell is lost in the process. Spatial transcriptomics (ST) is a recently developed technology and has the ability to map transcriptional signatures to distinct anatomical regions. To date, it has rarely been used in understanding lung tissues. In this first-in-class study, we employed a commercially available ST platform to investigate the spatial topography of gene expression of mouse lungs after immunization. To overcome the resolution limitation of current ST technology, we also applied state-of-art single-cell transcriptomics and single-cell epigenomics to jointly study the spatial localization, transcriptome, and epigenome of T cells induced by this immunization. The integrative analysis of three types of omics data provides an unprecedented and comprehensive way to examine the genomic basis and dynamics of lung immunization.

## RESULTS

### Cataloging 12 Lung Cell Types using scRNA-seq Data

In this study, we inoculated mice with heat-killed *K. pneumoniae* and assigned them into two groups, the immunized and the re-challenged groups. The immunized group was inoculated twice on day 0 and day 7, whereas the re-challenged group was additionally inoculated on day 13. On day 14, both groups of mice were sacrificed for tissue harvesting. Slices A1 and A2 were sectioned from the fresh frozen lung of the immunized mouse, whereas slices A3 and A4 were from the re-challenged mouse. In a separate cohort, lung tissue from the re-challenged mouse were harvested and single-cell suspension was obtained after enzymatic digestion. The four lung slices were used for spatial transcriptomics, and the single-cell suspension from the re-challenged mouse were subjected to scRNA-seq and scATAC-seq analyses. scRNA-seq data were integrated with scATAC-seq data via label transfer and were used as a reference to deconvolute spatial transcriptomics data (**Figures 1A**).

**Figure 1.**
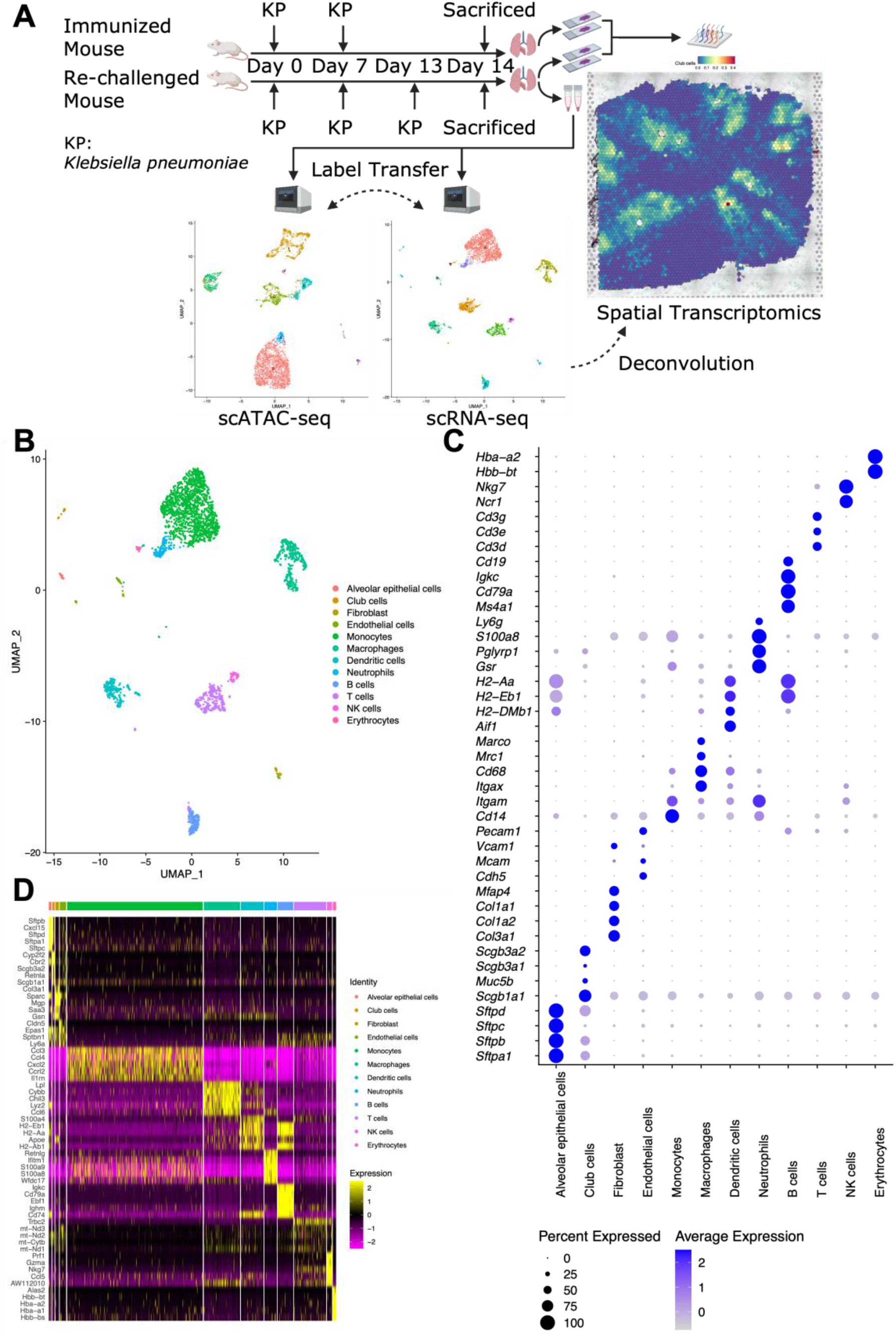
Generation of multi-omics datasets of mice lungs after immunization. (A) Overview of study design. scRNA-seq data work as a bridge to link scATAC-seq and spatial transcriptomics data. (B) UMAP plot of 12 lung cell types identified in scRNA-seq data, with manual annotation according to canonical markers. (C) Dot plot showing selected canonical markers for each cell type. (D) Heatmap showing top five (by log2-Fold Change) markers for each cell type.

We generated scRNA-seq data from 3,337 cells collected from the re-challenged mouse. Graph-based clustering identified ten clusters (**Figures S1A and S1C**). Cluster 1, which has an extremely high percentage of mitochondrial genes (**Figures S1B**), was excluded from downstream analyses. Cluster 6 consisted of several sporadic sub-clusters and was re-clustered into four clusters (**Figures S1D and S1E**). 2,834 cells with good quality were retained for the downstream analyses.

We carried out fine cluster annotation according to canonical markers and identified 12 lung cell types (**Figures 1B**), including alveolar epithelial cells, club cells, fibroblasts, endothelial cells, monocytes, macrophages, dendritic cells, neutrophils, B cells, T cells, NK cells, and erythrocytes. Selected canonical markers for each cell type were shown in the dot plot (**Figures 1C**, *e.g., Sftpb* for alveolar epithelial cells, *Scgb1a1* for club cells, and *Cd3d* for T cells). The top five (by log2-Fold Change) markers for each cell type were visualized using a heatmap (**Figures 1D**).

### Robust Cell-type Decomposition (RCTD) of Spatial Transcriptome using scRNA-seq Data

Although spatial transcriptomics provides additional spatial information, its resolution has not reached to single-cell level. Therefore, the expression profile of each spot in spatial transcriptomics is from a mixture of a few cells, typically one to ten. To better interpret spatial transcriptome data, it is vital to determine the proportions of different cell types within each spot. Using our finely annotated scRNA-seq data as a reference, we carried out deconvolution for each spot in the four slices using the robust cell-type decomposition (RCTD) method ^10^. The deconvolution results for slice A3 were shown as proportions of 12 cell types across slice A3 (**Figures S2A**).

To assess the robustness of this deconvolution method, we inspected whether well-characterized cell types were colocalized with the expression of their canonical markers, as well as corresponding histological structures. In slice A3, spots with a high proportion of club cells were around the histological airways and colocalized with spots with increased expression of *Scgb1a1* (**Figures 2Ai-iii**).

**Figure 2.**
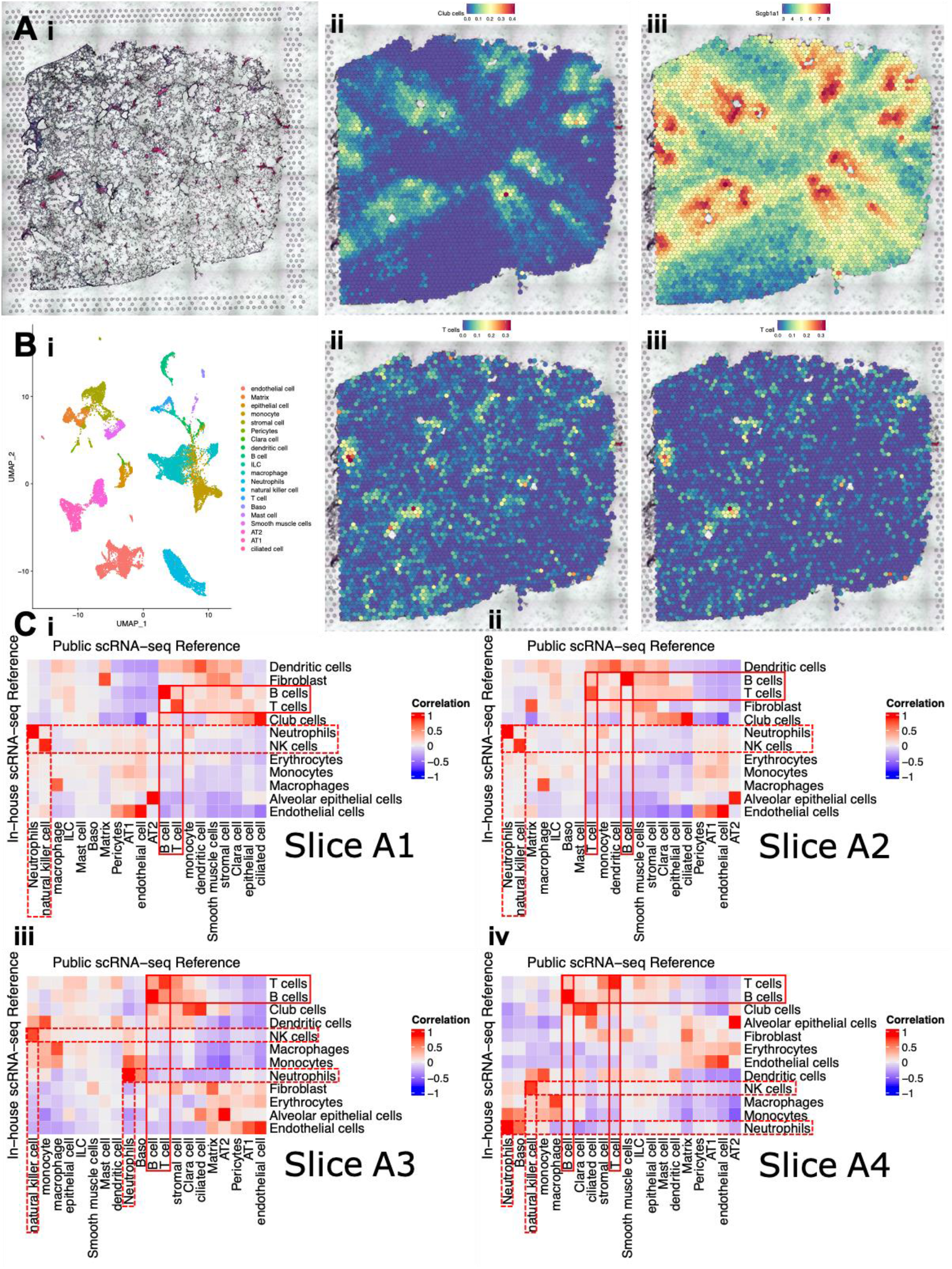
Validation of the robustness of deconvolution method for spatial transcriptomics. (A) Proportion of club cells colocalizing with the expression of *Scgb1a1* and the histological airways. (i) Histology of slice A3 showing the location of the airways. (ii) Proportion of club cells across slice A3, deconvoluted using in-house scRNA-seq data. (iii) Expression of *Scgb1a1*, a canonical marker for club cells, across slice A3. (B) Proportions of T cells deconvoluted using the two independent scRNA-seq references were quite similar. (i) UMAP plot of 20 lung cell types identified in Cohen *et al.’s* public scRNA-seq data (GSE119228). (ii) Proportion of T cells across slice A3, deconvoluted using in-house scRNA-seq data. (iii) Proportion of T cells across slice A3, deconvoluted using Cohen *et al.’s* public scRNA-seq data. (C)(i-iv) Correlation heatmap visualizing the proportions of cell types deconvoluted using inhouse (in rows, 12 types) and public (in columns, 20 types) scRNA-seq data were highly correlated in slice A1-A4. Pearson’s r values were indicated by the color bars. Red boxes with solid lines highlighted selected adaptive immune cells, T cells and B cells. Red boxes with dashed lines highlighted selected innate immune cells, neutrophils and NK cells.

To examine the robustness and impact of reference panel on deconvolution, another public scRNA-seq data (GEO: GSE119228) ^5^, including 20 annotated mouse lung cell types (**Figures 2Bi**), were also used as an independent reference to deconvolute spatial transcriptome for the four slices. The deconvolution results for slice A3, using public scRNA-seq reference, were shown (**Figures S2B**). In slice A3, proportions of T cells deconvoluted using in-house or public scRNA-seq references were quite similar (**Figures 2Bii and 2Biii**). Deconvolution results can also be represented using proportion matrices, whose rows indicate spots and columns indicate cell types. To quantify the similarity between the deconvolution results using the two independent references, Pearson correlation coefficients between the columns of the two proportion matrices were calculated and visualized using correlation heatmaps (**Figures 2Ci-iv**). Although we cannot perfectly match all the 12 cell types from our in-house data with the 20 cell types identified in this public dataset, we found the proportions of some adaptive immune cells (*e.g*., T cells and B cells) and innate immune cells (*e.g*., neutrophils and NK cells) were highly correlated with those deconvoluted using the other reference in all four slices. Other cell types were also highly correlated with their related cell types deconvoluted using the other reference (*e.g*., club cells in in-house data with ciliated cells in public data, alveolar epithelial cells in in-house data with AT2 in public data). These analyses validated the robustness of the robust cell-type decomposition (RCTD) method.

### Integrative Analysis of scRNA-seq and scATAC-seq Data Enabling the Identification of Th17 and Th1 Cells

In parallel with scRNA-seq, we also generated scATAC-seq data from 4,908 cells collected from the re-challenged mouse. After excluding cells with low quality, 4,794 cells were retained for downstream analyses. Leveraging our finely annotated scRNA-seq data, we identified the 12 lung cell types in scATAC-seq data via label transfer (**Figures 3A**). Proportions of the 12 cell types in scRNA-seq and scATAC-seq data were quite similar (**Figures 3B**), showing a good biological agreement between two types of data.

**Figure 3.**
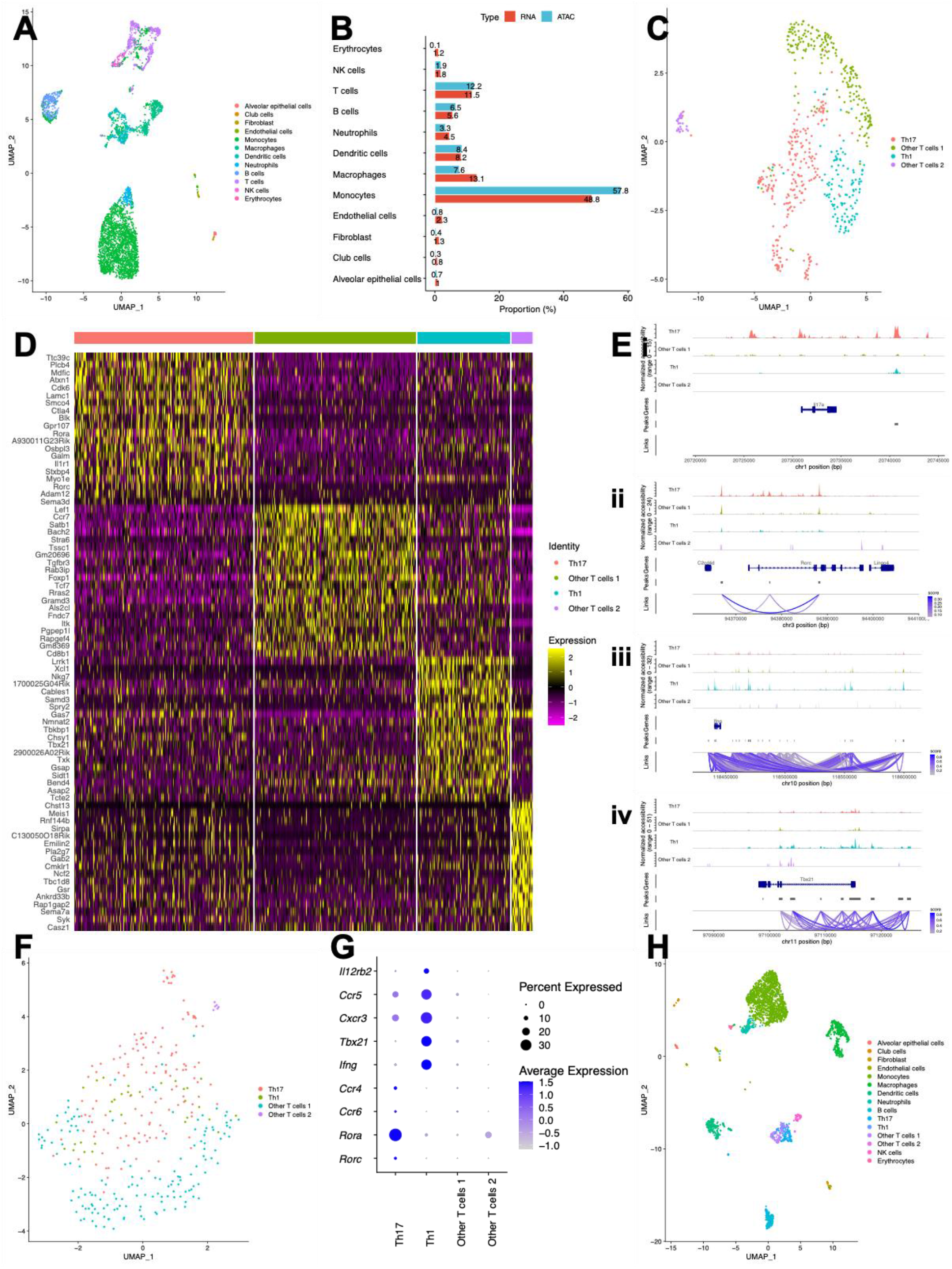
Identification of subtypes of T cells by integrating scRNA-seq and scATAC-seq data. (A) UMAP plot of 12 lung cell types in scATAC-seq data identified via label transfer. (B) Bar plot showing proportions of 12 cell types in scRNA-seq and scATAC-seq data were quite similar. (C) UMAP plot of four subtypes of T cells in scATAC-seq data. (D) Heatmap showing top 20 (by log2-Fold Change) markers (gene activity scores calculated using peaks) for each subtype of T cells in scATAC-seq data. (E)(i-iv) Peaks in genomic regions around *Il17a, Rorc, Ifng*, and *Tbx21*, canonical markers for Th17 and Th1 cells. (F) UMAP plot of four subtypes of T cells in scRNA-seq data identified via label transfer. (H) Dot plot showing selected canonical markers for Th17 and Th1 cells in scRNA-seq data. (G) UMAP plot of 15 lung cell types (including four subtypes of T cells) in scRNA-seq data.

Because we are interested in T cells in this study, we carried out re-clustering for T cells in scATAC-seq data and identified four subtypes of T cells (**Figures 3C**), including Th17 cells, Th1 cells, other T cells 1, and other T cells 2. The top 20 (by log2-Fold Change) markers (gene activity scores) for each subtype of T cells were visualized using a heatmap (**Figures 3D**). Canonical transcription factors for subtypes of T cells were shown on the heatmap (*e.g*., *Rorc* and *Rora* for Th17 cells, *Tbx21* for Th1 cells). *Cd8b1* was a marker for other T cells 1, suggesting they could be cytotoxic T cells.

To confirm the identity of Th17 and Th1 cells, further analyses were performed. For Th17 cells, there were much more peaks in genomic regions around *Il17a* and *Rorc*, compared with other subtypes of T cells (**Figures 3Ei and 3Eii**). In contrast, the peaks from Th1 cells dominated genomic regions around *Ifng* and *Tbx21* (**Figures 3Eiii and 3Eiv**). Motif footprinting analysis further confirmed RORC was dominated by Th17 cells, and TBX21 was dominated by Th1 cells (**Figures S3Ai-ii and S3Bi-ii**).

Although subtypes of T cells were almost indistinguishable only using scRNA-seq data, we managed to identify the four subtypes of T cells via label transfer from scATAC-seq data to scRNA-seq data (**Figures 3F**). To confirm the identity of Th17 and Th1 cells in scRNA-seq data, further analyses were performed. Selected canonical markers for Th17 and Th1 cells were shown in the dot plot (**Figures 3G**, *e.g*., *Rora* for Th17 cells, *Tbx21* for Th1 cells). The top 20 (by log2-Fold Change) markers for each subtype of T cells were visualized using a heatmap (**Figures S3C**). Canonical markers for subtypes of T cells can be seen in the heatmap (*e.g*., *Rora* for Th17 cells, *Ifng* for Th1 cells, and *Cd8b1* for other T cells 1). We also conducted gene regulatory network analysis using SCENIC ^11,12^ and identified RORA as a cell-type specific regulator for Th17 cells in scRNA-seq data (**Figures S3D**). By integrative analysis of scRNA-seq and scATAC-seq data, 15 lung cell types (including four subtypes of T cells) were identified in scRNA-seq data (**Figures 3H**), with the potential to be deconvoluted in spatial transcriptome.

### Dynamic Changes of Cell Locations upon *K. pneumoniae* Re-challenge

Using the updated scRNA-seq data with final 15 cell types, we carried out deconvolution again for each spot in the four slices (**Figures S4**). The proportions of 15 cell types across four slices were summarized by a box plot (**Figures 4A**). We performed *t*-tests to compare the differences between the immunized (slice A1 and A2) and the re-challenged mouse (slice A3 and A4), with significant differences found for each cell type. Generally, there were more monocytes, macrophages, dendritic cells, neutrophils, Th1 cells, and NK cells in the re-challenged mouse. In contrast, the immunized mouse had more B cells, Th17 cells, and other T cells 1 (**Figures 4A and S4**).

**Figure 4.**
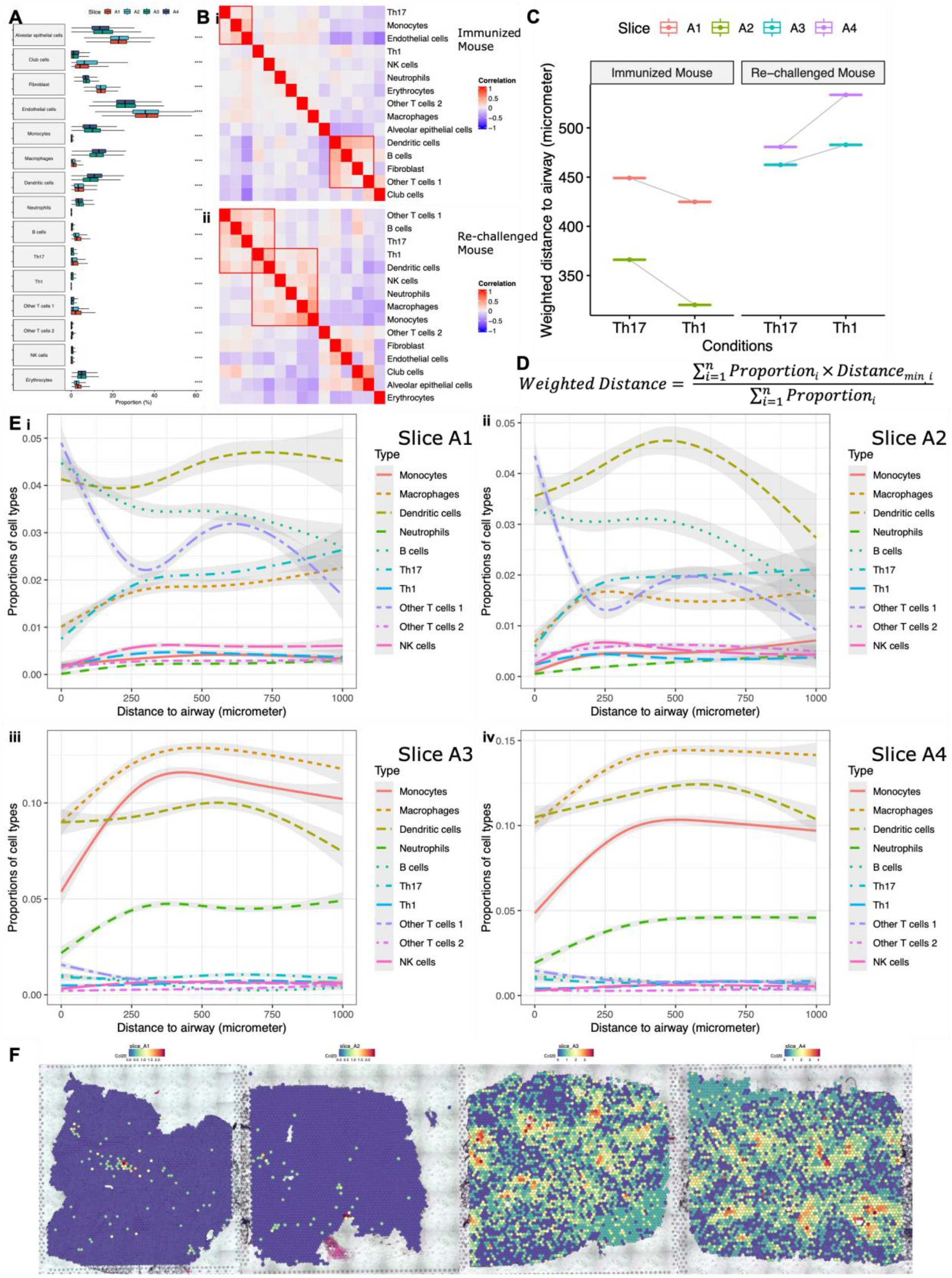
Spatial analyses of mice lungs after immunization. (A) Box plot showing proportions of 15 cell types across four slices were different. To compare the differences between the immunized (slice A1 and A2) and the re-challenged mouse (slice A3 and A4), *t*-tests were performed for each cell type. ****: p-value < 1×10^−4^. (B)(i-ii) Correlation heatmap visualizing the same-spot co-occurrence of 15 cell types in the immunized and the re-challenged mouse. ST spots from the slices for the immunized (slice A1 and A2) or the re-challenged mouse (slice A3 and A4) were pooled together, respectively. Red boxes highlighted cell types tending to appear in the same spots, that is to say, possibly close to one another *in vivo*. (C) Weighted distances to the airways for Th17 and Th1 cells in four slices, after excluding spots within the blood vessels and spots whose distances to the airways longer than 1,000 micrometers. (D) Formula defining weighted distance, allowing for each spot’s distance to the nearest airway and the proportions of Th17 and Th1 cells in each spot. (E)(i-iv) Proportions of immune cells over distance to the airways showing the spatial distribution of immune cells in four slices, after excluding spots within the blood vessels and spots whose distances to the airways longer than 1,000 micrometers. The curves were obtained from natural spline (with three degrees of freedom) regression. (F) Expression of *Ccl20*, a top distance-associated gene, across four slices.

After pooling spots from the slices for the immunized (slice A1 and A2) or the re-challenged mouse (slice A3 and A4), Pearson correlation coefficients between the columns of the proportion matrices were calculated. Significant same-spot co-occurrence of different cell types was found in the immunized and the re-challenged mouse (**Figures 4B**). For the immunized mouse, Th17 cells, monocytes, and endothelial cells tended to appear in the same spots. Dendritic cells, B cells, fibroblasts, and other T cells 1 were also close to one another. For the re-challenged mouse, other T cells 1, B cells, Th17 cells, Th1 cells, and dendritic cells often appeared in the same spots. While Th1 cells, NK cells, and myeloid cells were possibly close to one another *in vivo*. These observations remained unchanged with each slice analyzed separately (**Figures S5C**).

Localization and segmentation of airway and blood vessels are important in our analysis. We defined airways and blood vessels according to the proportion of club cells and the histological blood vessels (**Figures S5A and S5B**). After excluding spots within the blood vessels and spots whose distances to the airways longer than 1,000 micrometers, we calculated weighted distances to the airways for Th17 and Th1 cells in four slices, according to the formula (**Figures 4D**). Th17 cells were found closer to the airways than Th1 cells in the re-challenged mouse, whereas Th1 cells were closer to the airways in the immunized mouse (**Figures 4C**). This conclusion was independent of the definition of airways and blood vessels, since it remained unchanged even if a set of cut-offs were used to define these structures (**Figures S5D**).

To find the spatial distribution patterns of immune cells, natural spline regression was performed to fit the non-linear relationship between the proportions of immune cells and the distances to the airways. Generally, B cells and other T cells 1 were proximal to the airways in the immunized mouse, whereas neutrophils were distal to the airways in the re-challenged mouse (**Figures 4Ei-iv**). The same analysis for all 15 cell types was also performed (**Figures S5Ei-iv**). We also performed natural spline regression between the expression of genes and the distances to the airways. 3,655 distance-associated genes (FDR-adjusted P-value < 0.05 in both slices) were identified in the immunized mouse (**Table S1**), while 3,407 were identified in the re-challenged mouse (**Table S2**). Gene Ontology (GO) enrichment analysis was performed for these distance-associated genes ^13–15^. 1,293 significantly (FDR-adjusted P-value < 0.05) over-represented biological processes were identified for the immunized mouse, and 1,294 for the re-challenged mouse. The top 50 (hierarchically sorted by fold-enrichment) over-represented biological processes for the re-challenged mouse included many immune responses (*e.g*., proliferation, differentiation, activation, aggregation, adhesion, and chemotaxis of immune cells). In contrast, few immune responses were over-represented in the immunized mouse (**Table 1**). *Ccl20*, a top distance-associated gene, was highly expressed around the airways in the re-challenged mouse, instead of the immunized mouse (**Figures 4F**). Since CCL20 is capable of binding to CCR6, a chemokine receptor expressed on Th17 cells, this finding possibly explains why Th17 cells were closer to the airways than Th1 cells in the re-challenged mouse.

**Table 1.**
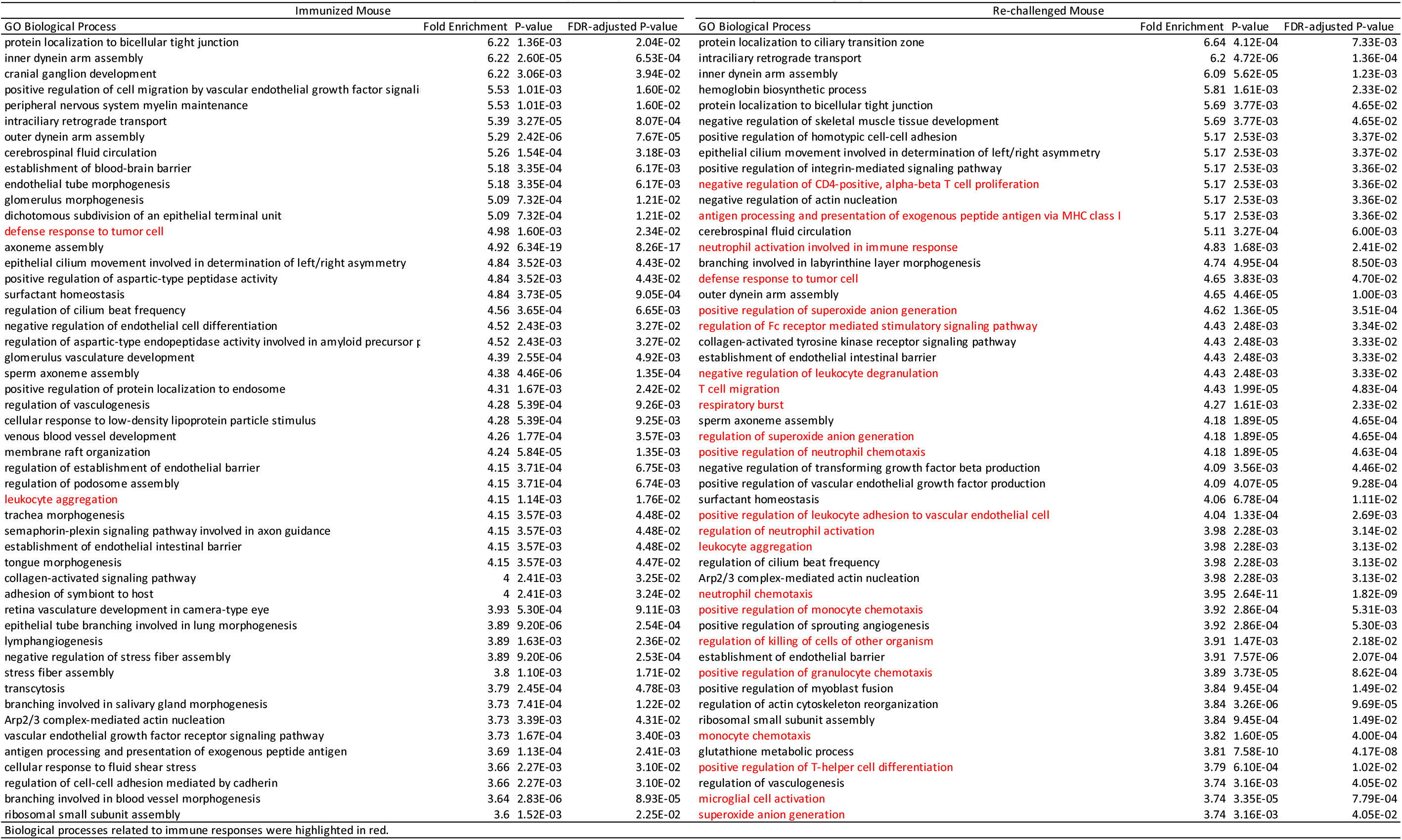
Top 50 (hierarchically sorted by fold-enrichment) over-represented biological processes for both mice

### Biological Differences upon *K. pneumoniae* Re-challenge

Upon *K. pneumoniae* re-challenge, the spatial transcriptome was tremendously changed (**Figures S6A**), while little batch effects were found between the two slices from the same mouse. The spots we defined as airways according to the proportion of club cells were also shown in a UMAP plot (**Figures S6B and S5A**), with significant biological differences found between the immunized and the re-challenged mouse.

Differential expression (DE) analysis was performed to compare the airways in the re-challenged mouse versus the immunized mouse, with 2,071 significantly (FDR-adjusted P-value < 0.05) differentially expressed genes (DEGs) identified (**Table S3**). To overcome the limitations of the analysis based on different thresholds of selected DEGs, we performed Gene Set Enrichment Analysis (GSEA) using the unfiltered, ranked gene list (including 16,937 genes), and found 379 significantly (FDR-adjusted P-value < 0.05) enriched GO biological processes. The top 30 (by normalized enrichment score) up-regulated and down-regulated pathways were shown by a lollipop plot (**Figures 5A**). Compared with the immunized mouse, most up-regulated pathways in the airways of the re-challenged mouse were related to immune responses (*e.g*., migration and chemotaxis of immune cells, and response to stimuli), whereas most down-regulated pathways were related to catabolic and metabolic processes, as well as ciliary functions. AW112010, a top DE gene up-regulated in the airways upon re-challenge, was highly expressed in the airways of the re-challenged mouse (**Figures 5B**). AW112010 has also been reported capable of promoting the differentiation of inflammatory T cells ^16^. *Cbr2*, a top DE gene down-regulated in the airways upon re-challenge, was highly expressed in the airways of the immunized mouse (**Figures 5C**), and may function in the metabolism of endogenous carbonyl compounds ^17^ and alveolar epithelial cell plasticity ^18^.

**Figure 5.**
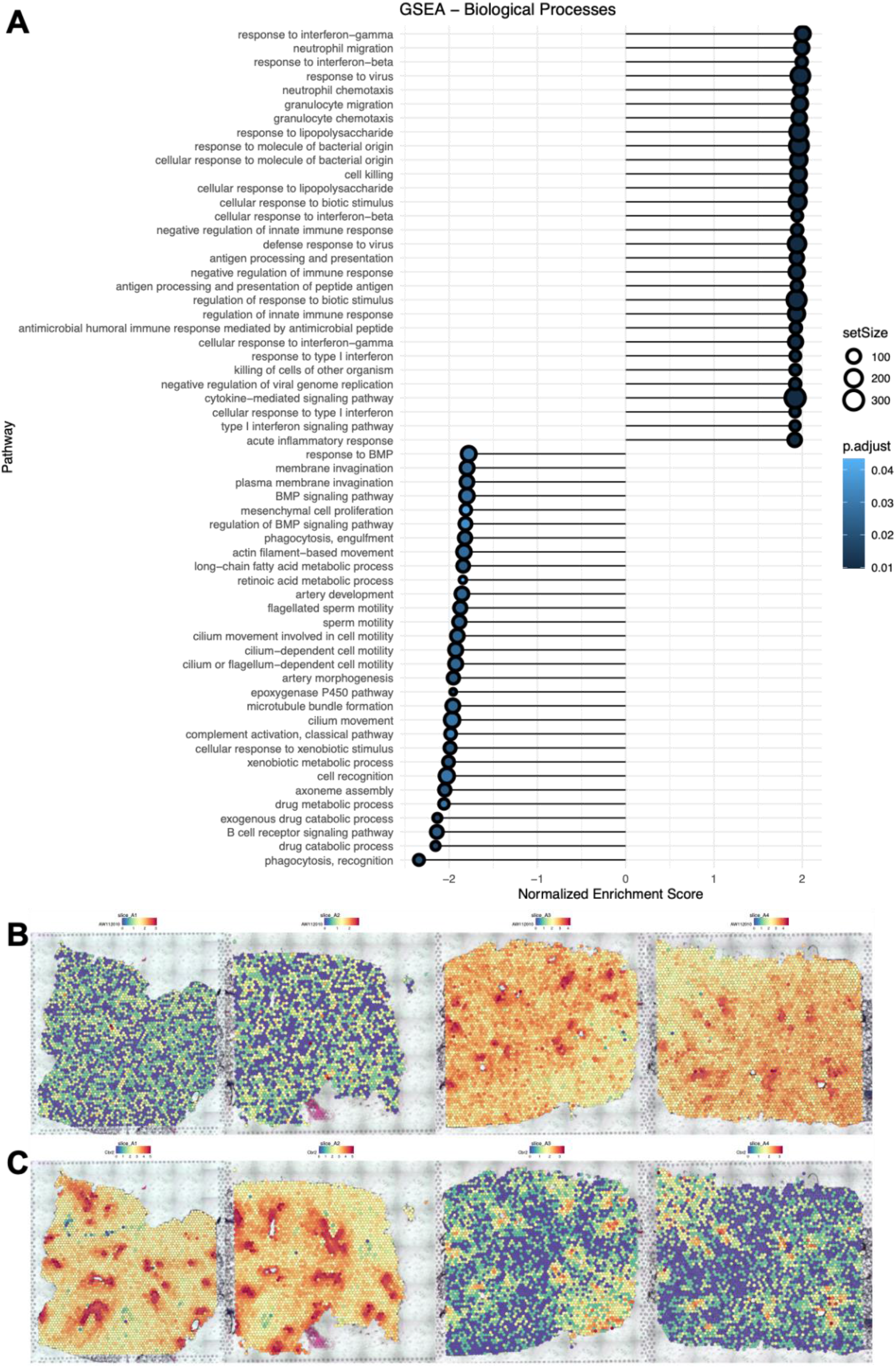
Differential expression analysis of spots annotated as airways in the immunized and the re-challenged mouse. (A) Gene Set Enrichment Analysis (GSEA) of all the 16,937 genes available in the DE analysis of the airways. (B) Expression of AW112010, a top DE gene up-regulated in the airways upon re-challenge, across four slices. (C) Expression of *Cbr2*, a top DE gene down-regulated in the airways upon re-challenge, across four slices.

### Spatial Transcriptomics Showing the Potential to Analyze Cell-cell Communication

In order to perform cell-cell communication networks analysis, cell-type enriched spots were identified according to their ranks of proportions of cell types (**Figures S6D**), with little difference found between the two slices from the same mouse (**Figures S6C**). Fibroblasts, endothelial cells, and erythrocytes were not included in the analysis due to the difficulty of interpreting the interactions with these cells.

Cell-cell communication networks among the cell-type enriched spots were inferred in each slice using CellChat (**Figures S6E**)^19^. To compare the differences of communication patterns between the re-challenged and the immunized mouse, differential interaction strength between cell-type enriched spots was shown by a heatmap (**Figures 6A**). The communication among myeloid cells, Th1 cells, and Th17 cells was increased in the re-challenged mouse, also revealed by our same-spot co-occurrence analysis. For the immunized mouse, dendritic cells could be the major sender of communication signals, while B cells and other T cells 1 were essential receivers of the signals.

**Figure 6.**
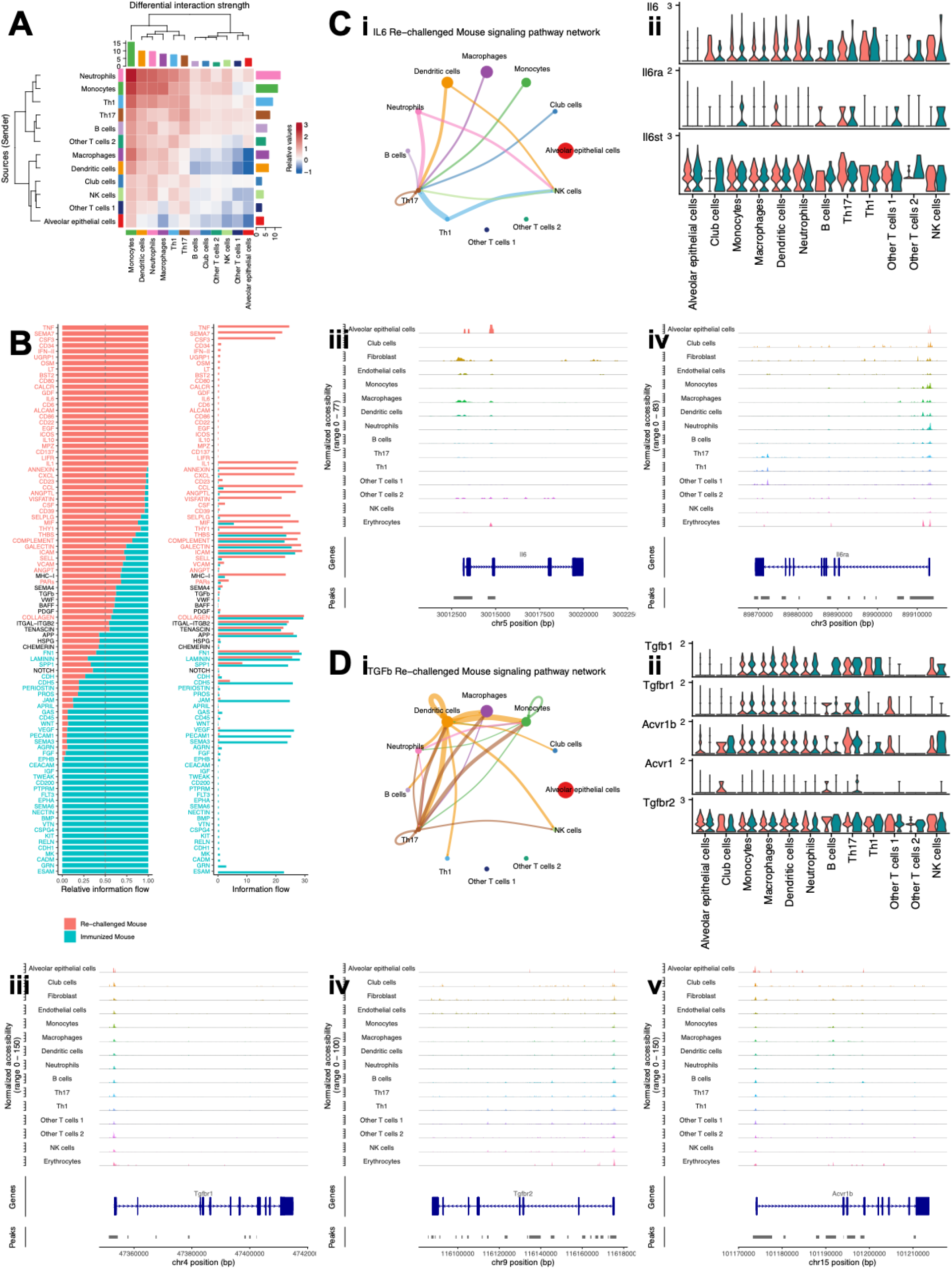
Cell-cell communication among cell-type enriched spots. (A) Heatmap showing differential interaction strength between cell-type enriched spots. Outgoing signals were shown in rows, while incoming signals were shown in columns. Increased (or decreased) signals in the re-challenged mouse compared to the immunized mouse were represented using red (or blue) in the color bar. The sum of values within the same column was summarized using the colored bar plot on the top. The sum of values within the same row was summarized using the colored bar plot on the right. (B) Bar plots showing overall information flow of each signaling pathway. Relative information flow was shown in the stacked bar plot, while raw information flow was shown in the regular bar plot. Enriched signaling pathways were colored in red or cyan. (C) Th17 enriched spots in the re-challenged mouse were the receivers of the IL6 signaling pathway. (i) Circle plot visualizing the inferred communication network of the IL6 signaling pathway in the re-challenged mouse. The network of the IL6 signaling pathway in the immunized mouse was not significant. Circle sizes represented the number of spots in each group. Edge colors were consistent with the senders of the signal (sources), and edge weights represented the interaction strength. (ii) Violin plot visualizing the expression of genes related to IL6 signaling pathway in cell-type enriched spots from the re-challenged mouse. Gene expression from slice A3 was colored in red, and that from slice A4 was colored in cyan. (iii-iv) Peaks in genomic regions around *Il6* and *Il6ra* in scATAC-seq data. (D) Th17 enriched spots in the re-challenged mouse were the receivers of the TGF-β signaling pathway. (i) Circle plot visualizing the inferred communication network of the TGF-β signaling pathway in the re-challenged mouse. The network of the TGF-β signaling pathway in the immunized mouse was also significant but not shown. Circle sizes represented the number of spots in each group. Edge colors were consistent with the senders of the signal (sources), and edge weights represented the interaction strength. (ii) Violin plot visualizing the expression of genes related to TGF-β signaling pathway in cell-type enriched spots from the re-challenged mouse. Gene expression from slice A3 was colored in red, and that from slice A4 was colored in cyan. (iii-v) Peaks in genomic regions around *Tgfbr1, Tgfbr2*, and *Acvr1b* in scATAC-seq data.

The conserved and context-specific signaling pathways were identified and visualized by bar plots (**Figures 6B**). TNF pathway was only turned on in the re-challenged mouse, while CXCL and CCL pathways were increased. We also confirmed some well-known signaling pathways in the cell-cell communication analysis of spatial transcriptome, outperforming the cell-cell communication analysis of scRNA-seq data. The performance of this analysis could be improved, as the resolution of spatial transcriptomics evolves. We found Th17 enriched spots in the re-challenged mouse were the receivers of the IL6 signaling pathway (**Figures 6Ci**), given the receptor gene *Il6ra* was highly expressed in Th17 enriched spots from slice A3 and slice A4 (**Figures 6Cii**). Leveraging scATAC-seq data, we confirmed macrophages and dendritic cells were the sources of the IL6 signaling (**Figures 6Ciii**), and Th17 cells were the targets (**Figures 6Civ**). For the TGF-β signaling pathway, we also found Th17 enriched spots in the re-challenged mouse were its receivers (**Figures 6Di**), with the receptor genes *Tgfbr1, Tgfbr2*, and *Acvr1b* found highly expressed in Th17 enriched spots from slice A3 and slice A4 (**Figures 6Dii**). By checking peaks in genomic regions around these receptor genes, Th17 cells were confirmed to have the capacity of expressing the receptors for the TGF-β signaling (**Figures 6Diii-v**). These data suggested that spatial transcriptomics data can be extrapolated and utilized to perform cell-cell communication networks analysis.

## DISCUSSION

As the development of single-cell sequencing technologies, scRNA-seq has been gradually applied to study immune cells and immune responses in mouse lungs ^5–8^. In many cases, T helper cells and cytotoxic T cells can be identified in these scRNA-seq studies, but not their subtypes. Few studies additionally utilize scATAC-seq to measure the chromatin architecture of immune cells in mouse lungs ^9^, which allows the identification of subtypes of T helper cells. As the birth of spatial transcriptomics, this cutting-edge technology is also adopted by this field to study mouse lungs infected by influenza ^20^.

To our knowledge, this is the first study to integrate spatial transcriptome with single-cell transcriptome and single-cell epigenome in mouse lungs. A key advantage of our study lies in the capture of intact anatomical structures of mouse lungs, providing location specific information and preserving information from cells prone to damage. Integrating with single-cell multi-omics profiled from matched tissue, sub-single-cell resolution were further achieved in our spatial analysis of cells residing in lung, especially TRM. In our study, we first identified 12 lung cell types in scRNA-seq data, covering major epithelial, mesenchymal, and immune cells. Using our finely annotated scRNA-seq data as a reference, we deconvoluted spatial transcriptome of the four slices and inferred proportions of cell types for each spot. We established an optimized deconvolution pipeline to accurately decipher specific cell-type composition at sub-single-cell resolution, by checking the correlation of deconvolution results using two independent references and the colocalization of canonical markers and histological structures. Integrating scATAC-seq data from the same cells processed side by side with scRNA-seq, we found epigenomic profiles provide more robust cell type identification, especially for lineage-specific T helper cells, and further identified four subtypes of T cells. Combining all three data modalities, we mapped the 15 lung cell types to histological structures for the four slices at sub-single-cell resolution.

Our data provides further insights into dynamic changes of cell locations upon *K. pneumoniae* re-challenge. We found Th17 cells were closer to the airways than Th1 cells in the re-challenged mouse, whereas Th1 cells were closer to the airways in the immunized mouse without re-challenge. This could be explained by the increased expression of *Ccl20* around the airways in the re-challenged mouse. *Ccl20*, a top distance-associated gene identified by our spatial analysis, has the ability to attract *Ccr6* expressing cells. The CCL20/CCR6 axis has been shown to play crucial roles in recruiting Th17 cells in many organs as well as various disease settings ^21,22^. We discovered different spatial distribution patterns of immune cells in the lungs of the two mice, finding B cells were proximal to the airways in the immunized mouse, whereas neutrophils were distal to the airways in the re-challenged mouse. We also identified thousands of distance-associated genes for the two mice by natural spline regression, confirming immune responses related genes were over-represented in the re-challenged mouse. These spatial analyses were only possible when location specific information was captured by spatial transcriptomics, highlighting the value of our study compared with conventional flow cytometric and transcriptomic approaches.

Comparing the biological differences in the airways upon *K. pneumoniae* re-challenge, we found pathways related to migration and chemotaxis of immune cells, and response to stimuli were up-regulated in the re-challenged mouse, whereas pathways related to catabolic and metabolic processes, and ciliary functions were up-regulated in the immunized mouse. We also performed cell-cell communication analysis of spatial transcriptome, providing evidence of lineage-specific T helper cells receiving designated cytokine signaling. Our study shows the potential to perform DE analysis between specific cells or regions across the slices and analyze cell-cell communication using spatial transcriptomics. The performance of these analyses could be improved as spatial transcriptomics advances towards single-cell resolution.

Several limitations were recognized in our study. First, we had a small sample size for our scRNA-seq and scATAC-seq data as the purpose of generating these data was to provide a reference for deconvolution, instead of carrying out a census of lung cell types. Second, there is no ground truth to evaluate deconvolution results, although we optimized our deconvolution pipeline by assessing its robustness. This issue could be eventually resolved as the resolution of the spatial transcriptomics technology improves. Third, our method to define airways and blood vessels may not be fully accurate and can be further improved by both high-resolution histological images and spatial transcriptome. Last, more biological replicates and experimental validations may be needed to extend this study. For example, the cellular source of CCL20 around the airways in the re-challenged mouse could be determined by immunofluorescence.

In summary, we presented a comprehensive single-cell multi-omics study on immunized mouse lungs and generated novel hypotheses for understanding underlying biological mechanism. Recently, spatial analysis has been made compatible with Formalin-Fixed Paraffin-Embedded (FFPE) tissue specimens. It is foreseeable that a massive amount of data will be generated from historically preserved samples. Our spatial transcriptomics data processing pipeline provides a timely solution to these analyses and contributes to advance the field of lung biology and respiratory medicine.

## MATERIALS AND METHODS

### Mouse models

All mice used in this study were wildtype and purchased from Jackson Lab (Cat# 000664). Animals were maintained in pathogen-free conditions in the animal facility at the University of Pittsburgh Medical Center. All experiments were approved by the University of Pittsburgh Institutional Animal Care and Use Committee.

### In vivo inflammation induction

6-8 weeks old C57BL/6 mice were immunized with heat-killed *K. Pneumoniae* (ATCC-43816) as previously described ^3^. Briefly, mice were injected with heat-killed *K. Pneumoniae* twice (Day0 and Day7) or three times (Day0, Day7, and Day13) intranasally and sacrificed on Day14. Lungs were removed and digested by Collagenase/DNase to obtain single-cell suspension. Mononuclear cells after red blood cell lysis and filtration with a 40 μM cell strainer were subjected to single cell RNA-seq (scRNA-seq) and single cell ATAC-seq (scATAC-seq) library prep following the protocols by 10x Genomics using the Chromium controller (10x Genomics). To yield sufficient IL-17A producing cells and reduce doublets formation, we targeted 3,000-5,000 cells/nuclei for recovery. Libraries were QC’ed on an Agilent TapeStation and sequenced on an Illumina Novaseq.

### Spatial Transcriptomics Experiment

We conducted ST experiment using 10X Genomics Visium platform.

#### Tissue harvesting

Mouse lungs were harvested, and the left lobes were inflated with 1mL mixture of 50% sterile PBS/ 50%Tissue-Tek OCT compound (SAKURA FINETEK) followed by frozen in alcohol bath on dry ice. OCT blocks were stored in −80C until further processing.

#### ST library prep

OCT blocks were sectioned at 10μm in thickness, 6.5mm X 6.5mm in size, attached to the Visium slides, then stained with hematoxylin and eosin following 10x Genomics Visium fresh frozen tissue processing protocol. H&E Images were taken by a fluorescence and tile scanning microscope (Olympus Fluoview 1000) then the slides underwent tissue removal and library generation per 10x Genomics demonstrated protocol.

### Raw Sequencing Data Processing

The sequenced scRNA-seq library was processed and aligned to mm10 mouse reference genome using Cell Ranger software (version 3.1.0) from 10x Genomics, with unique molecular identifier (UMI) counts summarized for each barcode. To distinguish cells from the background, cell calling was performed on the full raw UMI count matrix, with the filtered UMI count matrix generated (31,053 genes x 3,337 cells).

The sequenced scATAC-seq library was processed and aligned to mm10 mouse reference genome using Cell Ranger ATAC software (version 1.1.0) from 10x Genomics, with fragments and peak counts summarized for each barcode. To distinguish cells from the background, cell calling was performed on the full raw peak count matrix, with the filtered peak count matrix generated (84,317 peaks x 4,908 cells).

Each sequenced spatial transcriptomics library was processed and aligned to mm10 mouse reference genome using Space Ranger software (version 1.2.2) from 10x Genomics, with UMI counts summarized for each spot. To distinguish tissue overlaying spots from the background, tissue overlaying spots were detected according to the images. And only barcodes associated with these tissue overlaying spots were retained, with the filtered UMI count matrices generated. We also manually excluded spots not covered by tissue but not detected by Space Ranger and further filter the UMI count matrices (slice A1: 32,285 genes x 3,689 spots; slice A2: 32,285 genes x 2,840 spots; slice A3: 32,285 genes x 3,950 spots; slice A2: 32,285 genes x 3,765 spots).

### scRNA-seq Data Analysis

After imported into R, the filtered UMI count matrix was analyzed using the R package Seurat (version 4.0.1)^23^. The percentage of mitochondrial genes per cell was calculated for further check of the quality of cells. Regularized negative binomial regression (SCTransform)^24^ was used to normalize UMI count data, with the removal of confounding effects from mitochondrial mapping percentage. To improve the speed of the normalization, glmGamPoi ^25^ was invoked in the procedure. 3,000 highly variable genes were identified and used in principal component analysis to reduce dimensionality. We determined to use the first 50 principal components in clustering analysis according to the elbow plot. Uniform Manifold Approximation and Projection (UMAP) dimensionality reduction ^26^ was performed with the first 50 principal components as input to visualize cells. Using the Shared Nearest-neighbor (SNN) graph as input, cells were then clustered using the original Louvain algorithm with resolution = 0.2.

Cluster 1 was marked as low-quality cells and excluded from downstream analysis (2,834 cells retained) because its median percentage of mitochondrial genes was 87.7%, whereas those of all other clusters were lower than 10.5%. Markers for each cluster were identified using a Wilcoxon Rank Sum test with only.pos = TRUE, min.pct = 0.25, and logfc.threshold = 0.25. According to the UMAP plot, cluster 6 was found to consist of several sporadic sub-clusters. We isolated cells within cluster 6 and repeated the procedures from normalization to dimensionality reduction. These cells were then clustered using the original Louvain algorithm with resolution = 0.6. Markers were also identified as described above.

### Characterization of 12 Lung Cell Types in scRNA-seq Data

Fine cluster annotation was performed for the retained 12 clusters in scRNA-seq data according to canonical markers: alveolar epithelial cells (*Sftpa1, Sftpb, Sftpc*, and *Sftpd*), club cells (*Scgb1a1, Muc5b, Scgb3a1*, and *Scgb3a2*), fibroblasts (*Col3a1, Col1a2, Col1a1*, and *Mfap4*), endothelial cells (*Cdh5, Mcam, Vcam1*, and *Pecam1*), monocytes (*Cd14* and *Itgam*), macrophages (*Itgax, Cd68, Mrc1*, and *Marco*), dendritic cells (*Aif1, H2-DMb1, H2-Eb1*, and *H2-Aa*), neutrophils (*Gsr, Pglyrp1, S100a8*, and *Ly6g*), B cells (*Ms4a1, Cd79a, Igkc*, and *Cd19*), T cells (*Cd3d, Cd3e*, and *Cd3g*), NK cells (*Ncr1* and *Nkg7*), and erythrocytes (*Hbb-bt* and *Hba-a2*).

### Spatial Transcriptomics Data Analysis

After imported into R, the filtered UMI count matrix was analyzed using the R package Seurat (version 4.0.1)^23^. Regularized negative binomial regression (SCTransform)^24^ was used to normalize UMI count matrices, and glmGamPoi ^25^ was invoked in the procedure to improve the speed of the normalization. Four matrices from the four slices were merged to analyze them together. 3,000 highly variable genes were identified in each matrix, and the union set of them was set as highly variable genes for the merged data and used in principal component analysis to reduce dimensionality. We determined to use the first 30 principal components in clustering analysis according to the elbow plot. Uniform Manifold Approximation and Projection (UMAP) dimensionality reduction ^26^ was performed with the first 30 principal components as input to visualize spots.

### Deconvolution of Spatial Transcriptome

Deconvolution for each spot in the four slices was performed using the robust cell-type decomposition (RCTD) method ^10^. Before running the R package RCTD (version 1.2.0), scRNA-seq data used as a reference were processed, with gene expression matrix, the annotation for each cell, and the total UMI count for each cell extracted and saved in the RDS object. Spatial transcriptome for the four slices was also processed, with spot location matrices, gene expression matrices, and the total UMI count for each spot extracted and saved in the RDS object.

RCTD objects were created for each slice from the processed RDS objects, with max_cores = 24, test_mode = F, and CELL_MIN_INSTANCE = 6. RCTD pipeline was run on the RCTD objects, with doublet_mode = full. The deconvolution results were matrices of cell type weights for each spot. The cell type weights were normalized to make the sum of cell type weights in each spot equal to 1. Proportion matrices, whose rows indicate spots and columns indicate cell types, were then created and stored in the analyzed spatial transcriptome for loading.

In total, three scRNA-seq references were used in deconvolution, which were in-house scRNA-seq data with 12 cell types, in-house scRNA-seq data with 15 cell types (including four subtypes of T cells), and public scRNA-seq data with 20 cell types.

Pearson correlation coefficients between the columns (indicating different cell types) of the two proportion matrices deconvoluted using in-house (12 cell types) and public (20 cell types) references were calculated in each slice. Correlation r matrices were hierarchically clustered and visualized in heatmaps.

### scATAC-seq Data Analysis

After imported into R, the filtered peak count matrix was analyzed using the R package Signac (version 1.1.1)^27^. Gene annotations were extracted from Ensembl release 79 of the mm10 mouse reference genome. Nucleosome signal score and Transcriptional Start Site (TSS) enrichment score per cell were calculated for further check of the quality of cells. Cells whose fraction of fragments in peaks > 15, ratio of reads in genomic blacklist regions < 0.05, nucleosome signal score < 4, and TSS enrichment score > 2 were retained for downstream analysis. Latent Semantic Indexing (LSI)^28^, which is combined steps of Term Frequency–Inverse Document Frequency (TF-IDF) followed by Singular Value Decomposition (SVD), was used to normalize and reduce the dimensionality of peak count data, with all the peaks selected as variable features. We found the first LSI component was highly correlated with sequencing depth and determined to use the second to the fortieth LSI components in non-linear dimensionality reduction. Uniform Manifold Approximation and Projection (UMAP) dimensionality reduction ^26^ was performed with the second to the fortieth LSI components as input to visualize cells.

### Label Transferring from scRNA-seq Data to scATAC-seq Data

A gene activity matrix was created in scATAC-seq data by counting the number of fragments mapping to promoter or gene body regions of all protein-coding genes for each cell. Regularized negative binomial regression (SCTransform)^24^ was used to normalize the gene activity matrix. Transfer anchors were identified by canonical correlation analysis between the normalized gene activity matrix in scATAC-seq data and the normalized gene expression matrix in scRNA-seq data ^29^. Annotations were then transferred from scRNA-seq to scATAC-seq data with the second to the fortieth LSI components in scATAC-seq data used for weighting anchors. Canonical markers were checked in the gene activity matrix for each predicted cell type. The proportions of 12 cell types were also compared between scRNA-seq and scATAC-seq data.

### Characterization of Four Subtypes of T cells in scATAC-seq Data

We isolated predicted T cells in scATAC-seq data and repeated the procedures from normalization to non-linear dimensionality reduction. Using the Shared Nearest-neighbor (SNN) graph leveraging the second to the fortieth LSI components as input, T cells were then clustered into four subtypes using the Smart Local Movement (SLM) algorithm with resolution = 0.2.

Markers for each subtype were identified in the gene activity matrix using a Wilcoxon Rank Sum test with only.pos = TRUE, min.pct = 0.25, and logfc.threshold = 0.25. Differential accessible analysis was performed between the possible Th17 and Th1 cells using logistic regression, with fraction of fragments in peaks set as latent variable and min.pct = 0.1. Peaks in genomic regions around (including all the differentially accessible regions or 10,000 bps apart from the gene bodies) canonical markers for Th17 and Th1 cells were visualized. Motif footprinting analysis was also performed to provide supportive evidence. Taken together, four subtypes of T cells were annotated according to canonical markers: Th17 cells (*Il17a, Rorc*, and *Rora*), Th1 cells (*Ifng* and *Tbx21*), other T cells 1, and other T cells 2.

### Label Transferring from scATAC-seq Data to scRNA-seq Data

We isolated annotated T cells in scRNA-seq data and repeated the procedures from normalization to non-linear dimensionality reduction. Transfer anchors were identified by canonical correlation analysis between the normalized gene activity matrix of T cells in scATAC-seq data and the normalized gene expression matrix of T cells in scRNA-seq data ^29^. Annotations were then transferred from T cells in scATAC-seq to T cells in scRNA-seq data with the first 50 principal components in T cells from scRNA-seq data used for weighting anchors.

### Characterization of Four Subtypes of T cells in scATAC-seq Data

Canonical markers were checked for the predicted subtypes of T cells in scRNA-seq data (*Rorc*, *Rora, Ccr6*, and *Ccr4* for Th17 cells; *Ifng, Tbx21, Cxcr3, Ccr5*, and *Il12rb2* for Th1 cells). Markers for each predicted subtype of T cells were identified using a Wilcoxon Rank Sum test with only.pos = TRUE, min.pct = 0.25, and logfc.threshold = 0.25.

To detect active transcription factor (TF) modules, the R package SCENIC (version 1.2.4)^11,12^ was used to analyze the annotated scRNA-seq data, including 15 cell types. We downloaded the RcisTarget database containing transcription factor motif scores for gene promoter and around Transcription Start Site (TSS) for mm10 mouse reference genome from (https://resources.aertslab.org/cistarget/databases/mus_musculus/mm10/refseq_r80/mc9nr/gene_based/). The gene expression matrix was filtered according to default settings, and 9392 genes were retained and used to compute a gene-gene correlation matrix. Co-expression module detection was performed using the GENIE3 algorithm based on random forest. Transcription factor network analysis was performed to detect co-expression modules enriched for target genes of each candidate TF from the RcisTarget database, with regulons identified. The R package AUCell (version 1.12.0) was used to compute a score for each TF module in each cell.

Regulon Specificity Score (RSS) was calculated for regulons in each cell type according to Area Under the Curve (AUC) of regulons. Cell-type specific regulators were then identified, and those for the four subtypes of T cells were visualized to provide supportive evidence (Rora_extended (15g) for Th17 cells).

### Same-spot Co-occurrence Analysis

Pearson correlation coefficients between the columns (indicating different cell types) of the proportion matrices were calculated in each slice to broadly assess spatial cell type co-occurrence in the same spot. Spots from the slices for the immunized (slice A1 and A2) or the re-challenged mouse (slice A3 and A4) were pooled together, respectively, and Pearson correlation coefficients were also calculated for them. Correlation r matrices were hierarchically clustered and visualized in heatmaps.

### Defining Airways and Blood Vessels in Four Slices

Airways were defined according to the proportion of club cells in four slices. We manually set the thresholds in each slice to match the selected spots with the histological airways. Spots whose proportion of club cells higher than the thresholds were defined as airways (20% for slice A1, 20% for slice A2, 10% for slice A3, and 10% for slice A4).

Blood vessels were defined according to the histological blood vessels. We created a training set using manual annotation of histological structures in the image of slice A1 and trained a random trees pixel classifier using QuPath (version 0.2.3)^30^ with downsample = 16. The probability of blood vessels was predicted for each pixel in the image of four slices using the trained classifier. If the probability of blood vessels in the spot corresponding pixel was higher than 0.5, the spot would be defined as blood vessels.

Alternatively, the proportion of club cells and the expression of *Mgp* were used to define airways and blood vessels with different cut-offs, from 90^th^ quantile to 95^th^ quantile. Spots whose proportion or expression higher than the selected quantile were defined as airways or blood vessels.

### Spatial Transcriptomics Spot Distance-based Analyses

For all distance-based analyses, spots defined as blood vessels and spots whose distances to the airways longer than 1,000 micrometers were excluded. Because cells within the blood vessels were different from those in the lung parenchyma, and the analyses including cells extremely distal to the airways were not stable, given the capture area of each slice was only 6.5 × 6.5 mm^2^. Only a few spots’ distances to the airways were longer than 1,000 micrometers (5.89% for slice A1, 5.75% for slice A2, 15.00% for slice A3, and 11.43% for slice A4).

Weighted distances to the airways for Th17 and Th1 cells were calculated in four slices according to the formula, allowing for each spot’s distance to the nearest airway and the proportions of Th17 and Th1 cells in each spot.

To find the spatial distribution patterns of cell types, natural spline regression (with three degrees of freedom) was performed to fit the non-linear relationship between the proportions of cell types and the distances to the airways.

To identify distance-associated genes, we constructed natural splines (with three degrees of freedom) for the distances to the airways in each slice and created design matrices. We then fitted linear models for each gene in normalized gene expression matrices, with the R package limma (version 3.46.0)^31^ invoked to speed up the procedure. P-values for each spline in each slice were corrected for multiple testing using Benjamini-Hochberg correction. Genes whose FDR-adjusted P-values < 0.05 for at least one spline were considered significant in that slice. And genes significant in both slices for the immunized (slice A1 and A2) or the re-challenged mouse (slice A3 and A4) were defined as distance-associated genes.

### Gene Ontology (GO) Enrichment Analysis

Gene ontology enrichment analysis was performed on the distance-associated genes identified in the immunized and re-challenged mouse using PANTHER Classification System (version 16.0)^13–15^. GO biological process complete was used as annotation dataset, and the analysis was performed using Fisher’s exact test, with FDR adjustment for multiple testing. Over-represented pathways were then hierarchically clustered.

### Differential Expression Analysis

Differential expression analysis was performed to compare the airways in the re-challenged mouse versus the immunized mouse using the R package MAST (version 1.16.0)^32^. MAST procedure was invoked in Seurat FindMarkers function (test.use = MAST) with logfc.threshold = 0, min.pct = 0, in order to obtain an unfiltered gene list. The gene list was then ranked by log2-fold change (L2FC). Genes whose L2FC equal to 0 were excluded due to their ranks were not available.

### Gene Set Enrichment Analysis (GSEA)

To overcome the limitations of the analysis based on manually selected DEGs, Gene Set Enrichment Analysis (GSEA)^33^ was performed on the unfiltered, ranked gene list (including 16,937 genes) using the R package clusterProfiler (version 3.18.1)^34^. The parameters for the clusterProfiler gseGO function were set as ont = BP, keyType = SYMBOL, nPerm = 10,000, minGSSize = 3, maxGSSize = 800, pvalueCutoff = 0.05, OrgDb = org.Mm.eg.db, pAdjustMethod = fdr. The top 30 (by normalized enrichment score) up-regulated and down-regulated pathways were then visualized by a lollipop plot.

### Defining Cell-type Enriched Spot in Spatial Transcriptomics Data

The expression profile of each spot in spatial transcriptomics is a mixture of a few cells, and it is irrational to annotate a spot with a cell type directly. To perform cell-cell communication analysis of spatial transcriptome, we annotated spots as cell-type enriched spots according to their proportions of cell types.

The mean proportions of cell types were available for each slice, as well as the number of spots for each slice. The mean proportions could be interpreted as the expected proportions of cell types in each spot. The product of the mean proportions and the number of spots could be interpreted as how many spots could represent each cell type on average. The spots with the highest proportions were selected according to the product and defined as cell-type enriched spots. Spots defined as cell-type enriched spots for multiple cell types were then excluded.

For example, the mean proportion of club cells was 6.39% in slice A1, and the number of spots was 3,689 in slice A1. Thus, the top 236 spots with the highest proportion of club cells were defined as club cell enriched spots. A spot would be excluded if the spot was defined as a club cell enriched spot and a Th17 enriched spot simultaneously.

### Cell-cell Communication Analysis of Spatial Transcriptome

Cell-cell communication analysis was performed using CellChat ^19^. Before running the R package CellChat (version 1.1.0), spatial transcriptome for the four slices was processed, with normalized gene expression matrices, the annotations for cell-type enriched spots extracted and saved in the CellChat object. The processed data from both slices for the immunized (slice A1 and A2) or the re-challenged mouse were also pooled together and saved in the CellChat objects to compare the differences between the two mice.

A manually curated database of literature-supported ligand-receptor interactions in mouse was loaded for the analysis. And all ligand-receptor interactions, including paracrine/autocrine signaling interactions, extracellular matrix (ECM)-receptor interactions, and cell-cell contact interactions, were included in the analysis.

Over-expressed ligands or receptors were first identified for each type of cell-type enriched spots in each CellChat object. Over-expressed ligand-receptor interactions were also identified if either ligand or receptor was over-expressed. Then, we computed the communication probability and inferred cellular communication network according to default settings. The communication probability at signaling pathway level was computed by summarizing the communication probabilities of all ligands-receptors interactions associated with each signaling pathway. The aggregated cell-cell communication network was calculated by counting the number of links or summarizing the communication probability and visualized in four slices using heatmaps.

To figure out in which type of cell-type enriched spots interactions significantly changed in the re-challenged mouse versus the immunized mouse, differential interaction strength was identified and visualized using a heatmap. The conserved and context-specific signaling pathways were identified by comparing the information flow for each signaling pathway, which was defined by the total weights in the network. Selected signaling pathways were visualized by circle plots using CellChat netVisual_aggregate function.

## Supporting information

Figures with high resolution

## Acknowledgements

This research was supported in part by the University of Pittsburgh Center for Research Computing through the resources provided.

## Author contributions

Z.X., W.C., and K.C. conceived the project and designed the experiments. L.F. and F.W. performed scRNA-seq, scATAC-seq, and ST experiments. Z.X. and X.W. performed scRNA-seq, scATAC-seq, and ST data analysis. J.W. provided guidance for deconvolution of ST data. Z.X., W.C., and K.C. wrote the manuscript with input from all authors. W.C. and K.C. supervised the work.

## Compliance with ethics guidelines

None of authors have any conflict of interest to report. All animal protocols and procedures were reviewed and approved by the University of Pittsburgh Institutional Animal Care and Use Committee.

## Data and materials availability

Raw and processed data of scRNA-seq, scATAC-seq and ST will be deposited to Gene Expression Omnibus (GEO) upon acceptance of the paper. Code and scripts necessary to repeat analyses in this manuscript are available upon request.

