## Supplemental Figures for "Integrative Analysis of Spatial Transcriptome with Single-cell Transcriptome and Single-cell Epigenome in Mouse Lungs after Immunization"

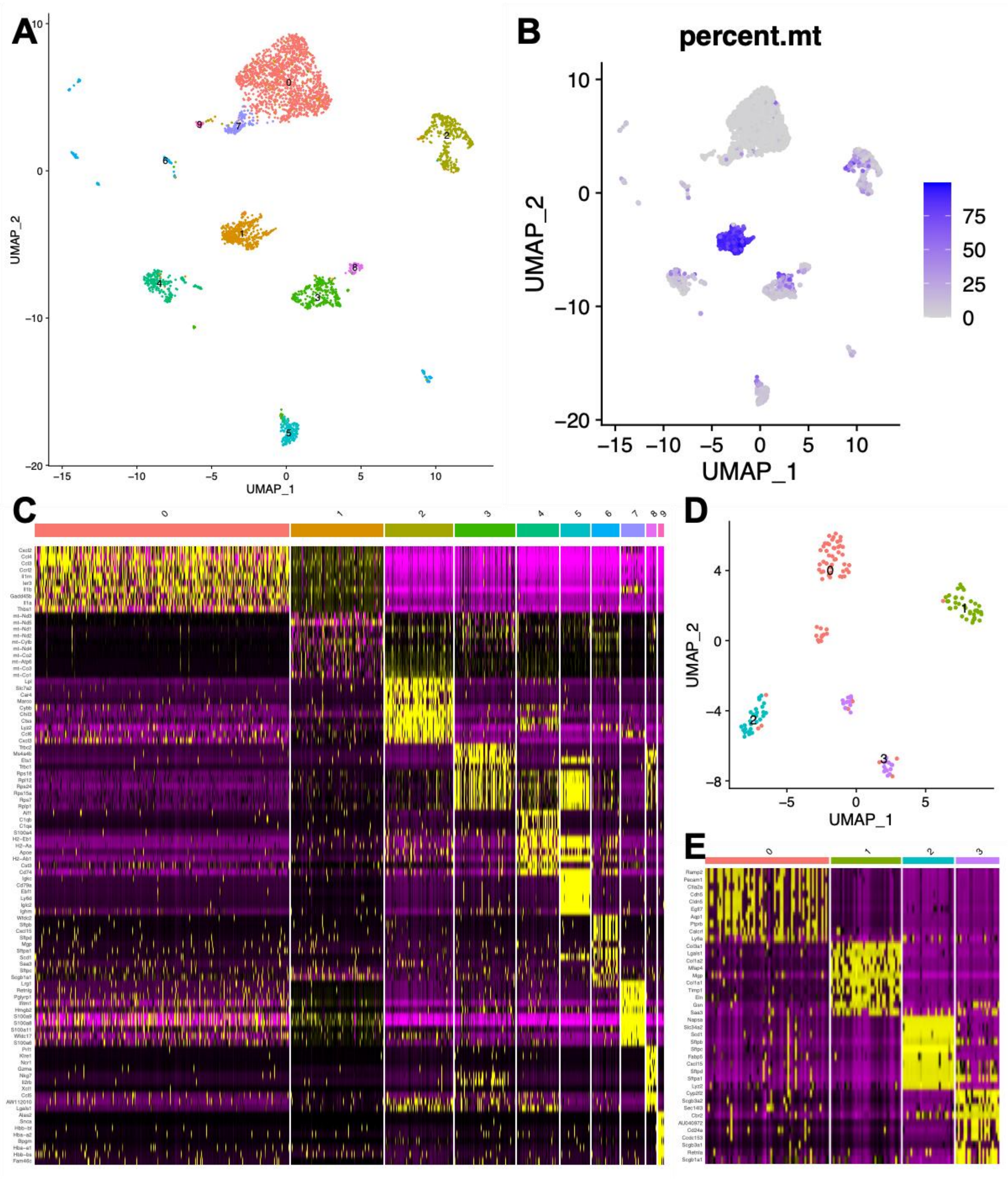


**Figure S1. Graph-based clustering of scRNA-seq data, related to Figure 1**

(A) UMAP plot of ten clusters identified in scRNA-seq data.

(B) Distribution of cells with a high percentage of mitochondrial genes. Cluster 1, which has an extremely high percentage of mitochondrial genes, was excluded from downstream analyses.

(C) Heatmap showing top ten (by log2-Fold Change) markers for each cluster.

(D) UMAP plot of four sub-clusters identified in cluster 6.

(E) Heatmap showing top ten (by log2-Fold Change) markers for each sub-cluster identified in cluster 6.

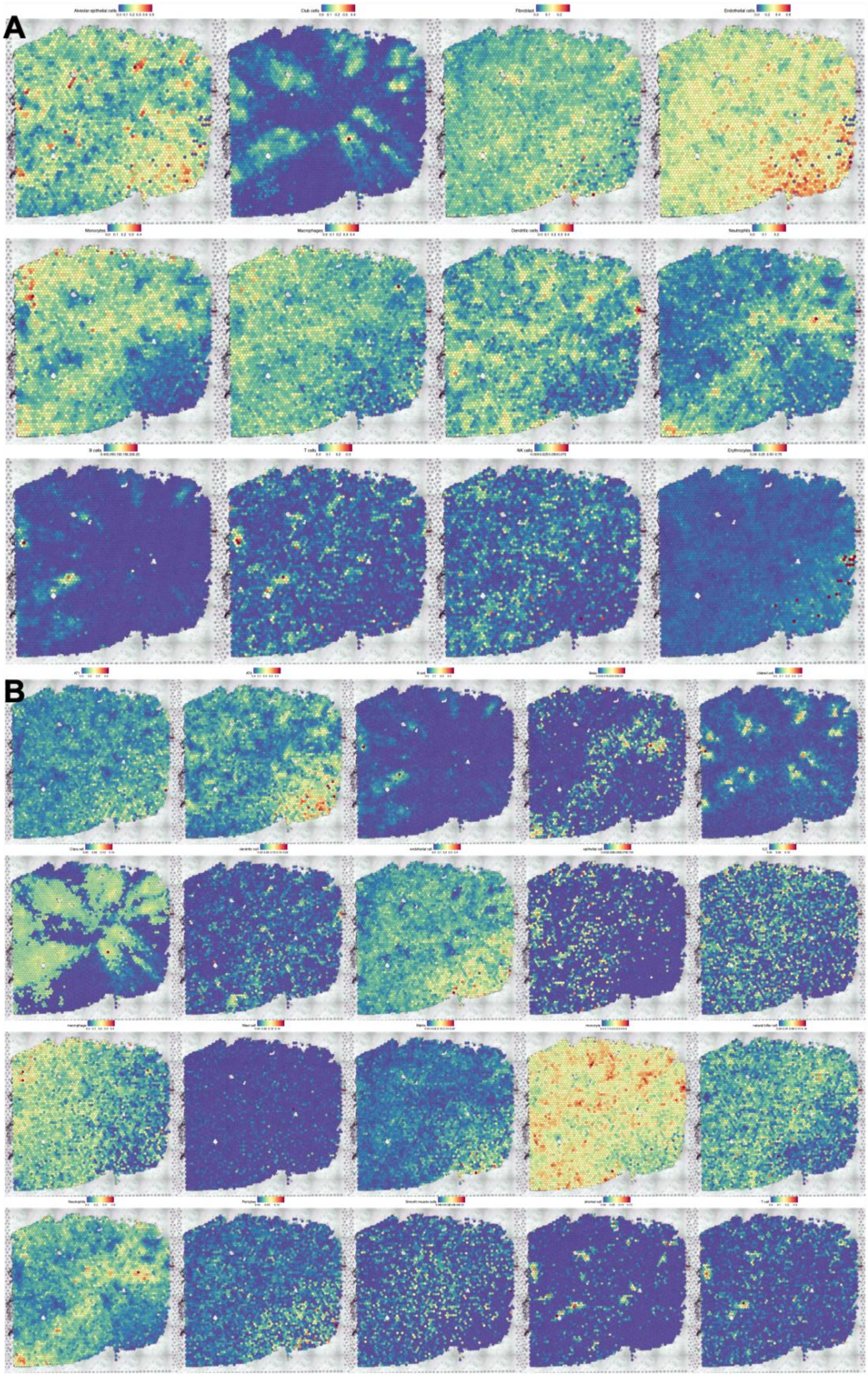
A
Alveolar epithelial cells
Club cells
Fibroblast
Endothelial cells
Monocytes
Macrophages
Dendritic cells
Neutrophils
B cells
T cells
NK cells
Erythrocytes
B
AT1
AT2
B cell
ciliated cell
Clara cell
dendritic cell
endothelial cell
macrophage
Mast cell
monocyte
natural killer cell
Neutrophils
Pericytes
Smooth muscle cells
stromal cell
T cell

**Figure S2. Proportions of cell types deconvoluted using two independent scRNA-seq references, related to Figure 2**

(A) Proportions of 12 cell types across slice A3, deconvoluted using in-house scRNA-seq data.

(B) Proportions of 20 cell types across slice A3, deconvoluted using Cohen *et al*.'s public scRNA-seq data.

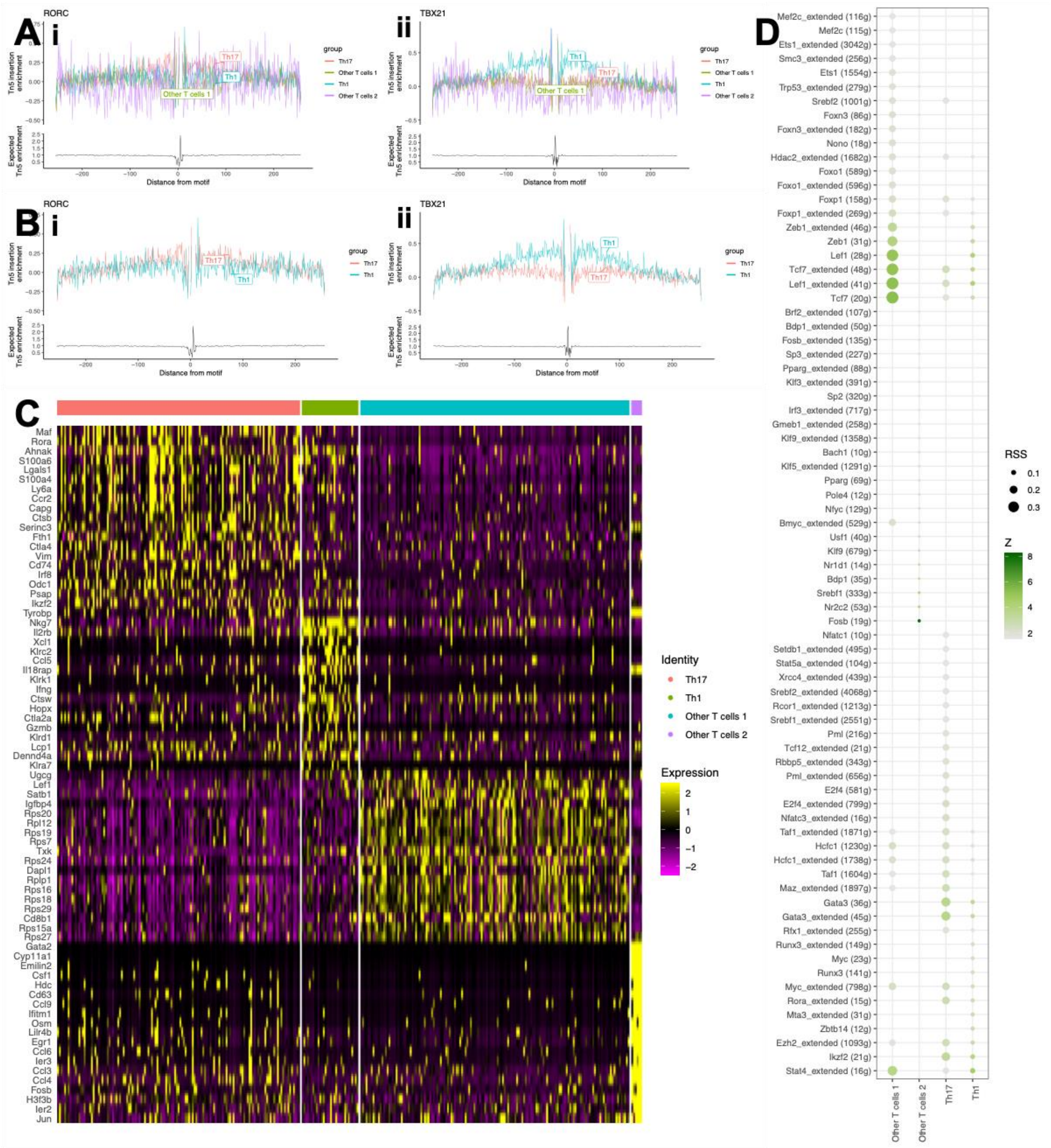

**Figure S3. Validation of the subtypes of T cells, related to Figure 3**

(A)(i-ii) Motif footprinting of RORC and TBX21, canonical markers for Th17 and Th1 cells, across four subtypes of T cells.

(B)(i-ii) Motif footprinting of RORC and TBX21, canonical markers for Th17 and Th1 cells, across Th17 and Th1 cells.

(C) Heatmap showing top 20 (by log2-Fold Change) markers for each subtype of T cells in scRNA-seq data.

(D) Dot plot showing cell-type specific regulators identified using SCENIC for each subtype of T cells in scRNA-seq data.

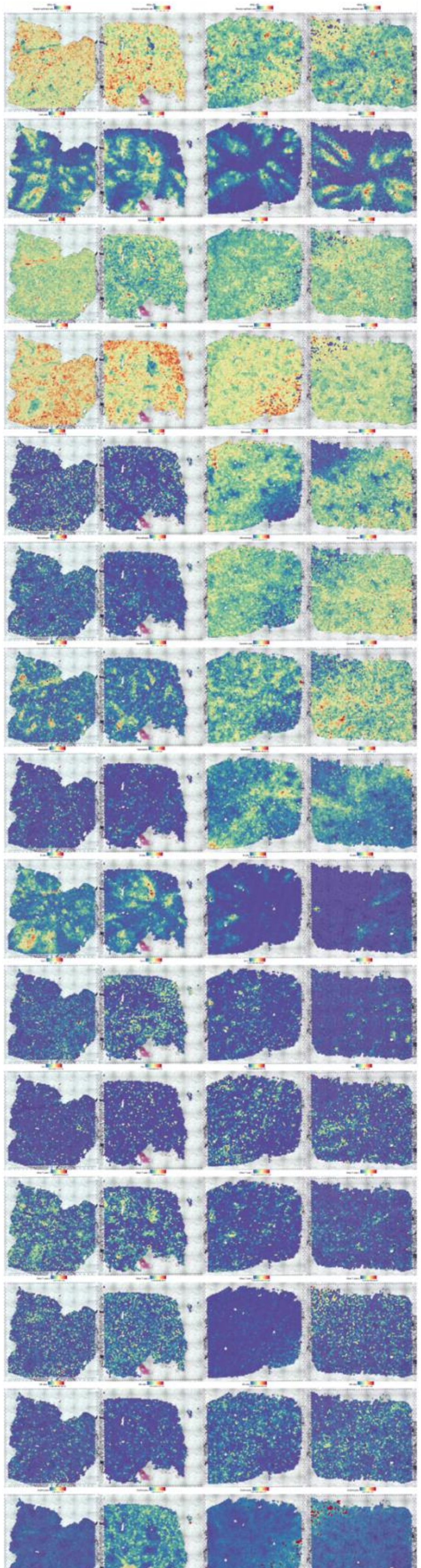

**Figure S4. Proportions of cell types across four slices, related to Figure 4**

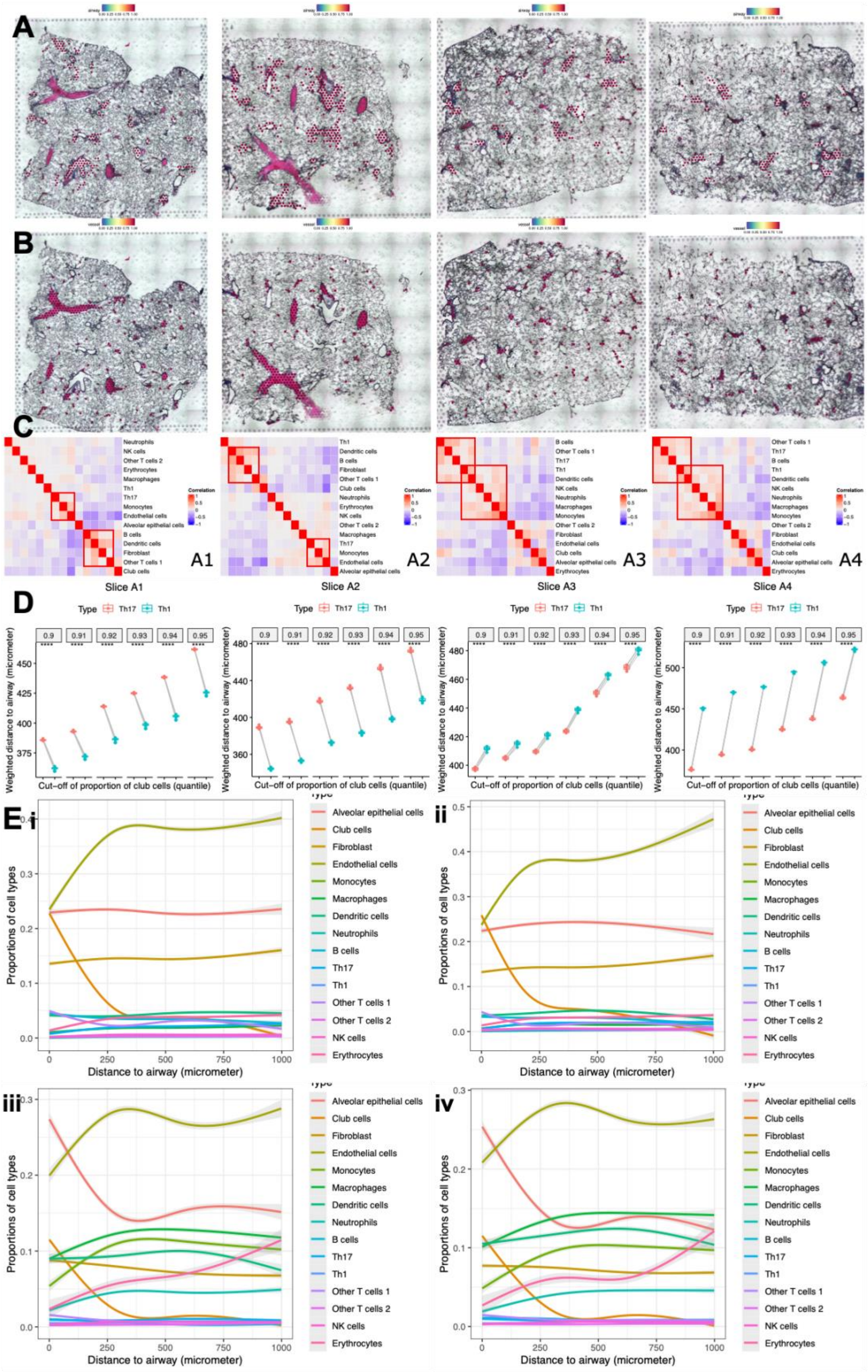
A
B
C
D
E i
ii
iii
iv
A1
A2
A3
A4
Slice A1
Slice A2
Slice A3
Slice A4
Type Th17 Th1
0.9
0.91
0.92
0.93
0.94
0.95
Weighted distance to airway (micrometer)
Cut-off of proportion of club cells (quantile)
Proportions of cell types
Distance to airway (micrometer)
Alveolar epithelial cells
Club cells
Fibroblast
Endothelial cells
Monocytes
Macrophages
Dendritic cells
Neutrophils
B cells
Th17
Th1
Other T cells 1
Other T cells 2
NK cells
Erythrocytes

**Figure S5. Spatial analyses of mice lungs after immunization, related to Figure 4**

(A) ST spots annotated as airways across four slices.

(B) ST spots annotated as blood vessels across four slices.

(C) Correlation heatmap visualizing the same-spot co-occurrence of 15 cell types in four slices. Red boxes highlighted cell types tending to appear in the same spots, that is to say, possibly close to one another *in vivo*.

(D) Box plots showing weighted distances to the airways for Th17 and Th1 in four slices, after excluding spots within the blood vessels and spots whose distances to the airways longer than 1,000 micrometers. The proportion of club cells and the expression of *Mgp* were used to define airways and blood vessels with different cut-offs, from $90^{th}$ quantile to $95^{th}$ quantile. *t*-tests were performed for each cut-off to define airway. $^{ns}$: not significant; $^{***}$: p-value < 0.001; $^{****}$: p-value < $1x10^{-4}$.

(E)(i-iv) Proportions of cell types over distance to airway showing the spatial distribution of cell types in four slices, after excluding spots within the blood vessels and spots whose distances to the airways longer than 1,000 micrometers. The curves were obtained from natural spline (with three degrees of freedom) regression.

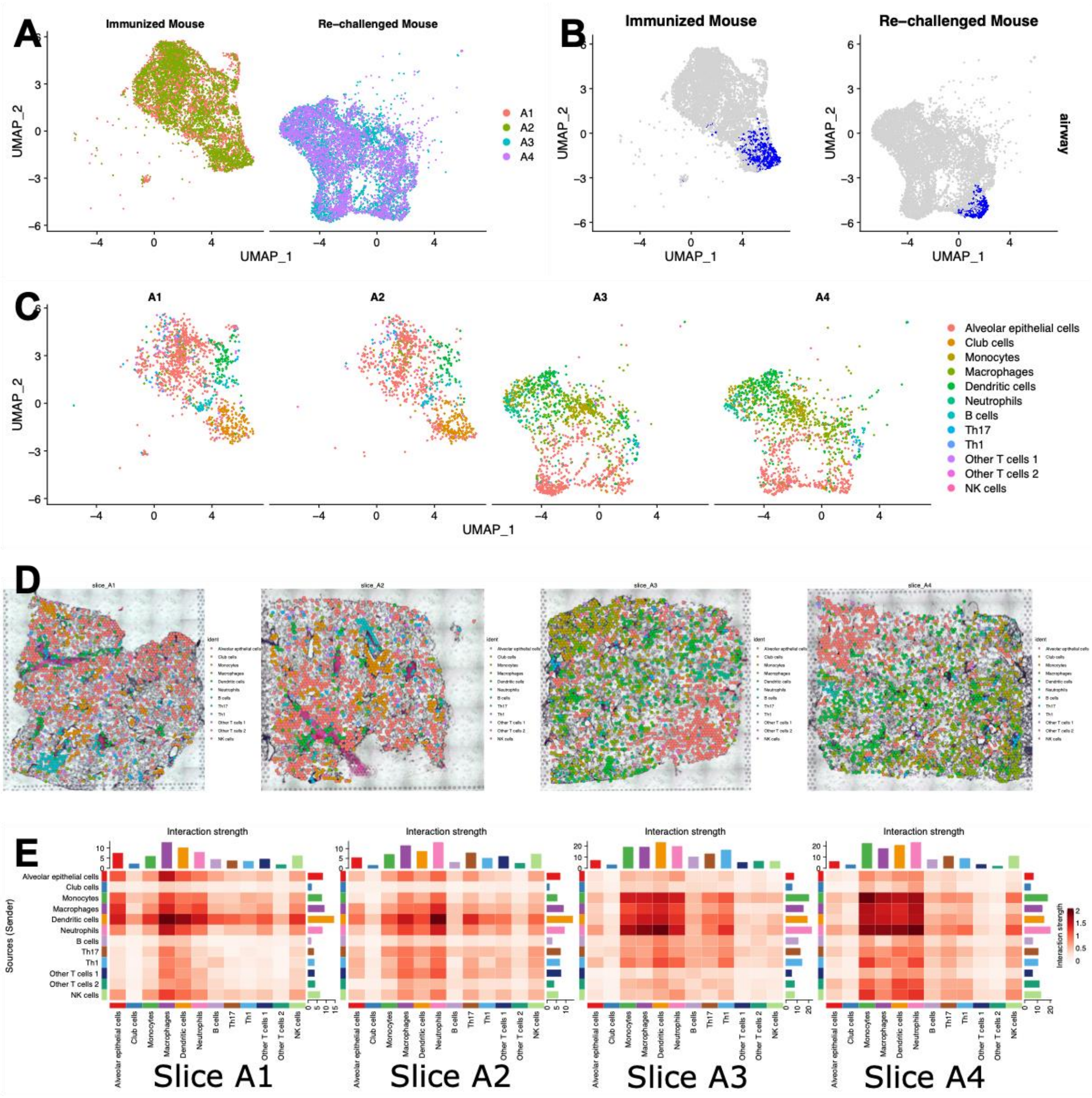


**Figure S6. Defining cell-type enriched spots, related to Figure 5 and 6**

(A) UMAP plot showing considerable biological differences between the immunized and the re-challenged mouse.

(B) UMAP plot showing spots annotated as airways in the immunized and the re-challenged mouse.

(C) UMAP plot showing spots annotated as cell-type enriched spots in four slices.

(D) Cell-type enriched spots across four slices.

(E) Heatmap showing interaction strength in four slices. Outgoing signals were shown in rows, while incoming signals were shown in columns. Interaction strength was represented in the color bar. The sum of values within the same column was summarized using the colored bar plot on the top. The sum of values within the same row was summarized using the colored bar plot on the right.
