## Supplementary figures and images for "Integrative Analysis of Spatial Transcriptome with Single-cell Transcriptome and Single-cell Epigenome in Mouse Lungs after Immunization"

### ST_figure_1.pdf

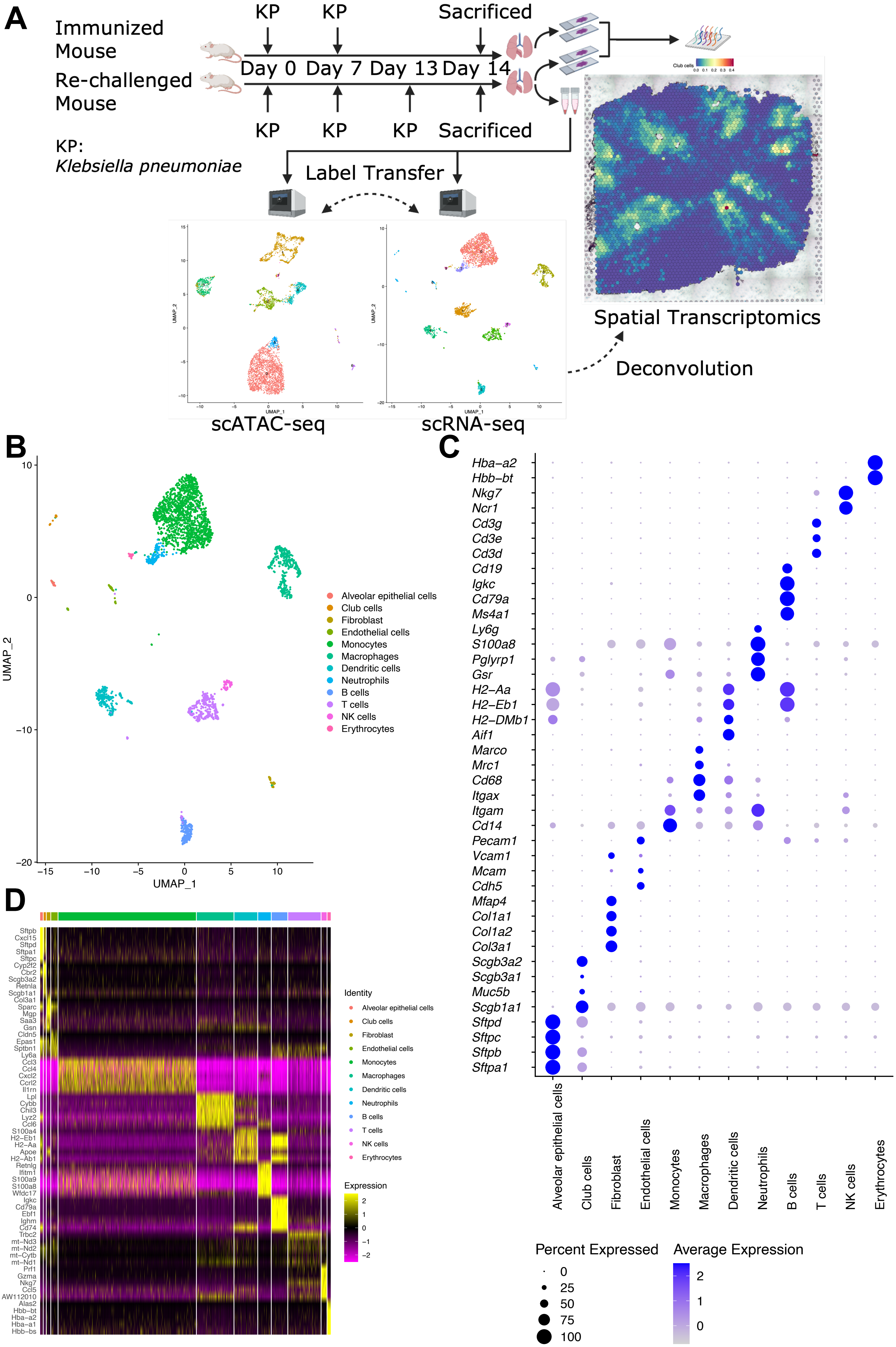

### ST_figure_2.pdf

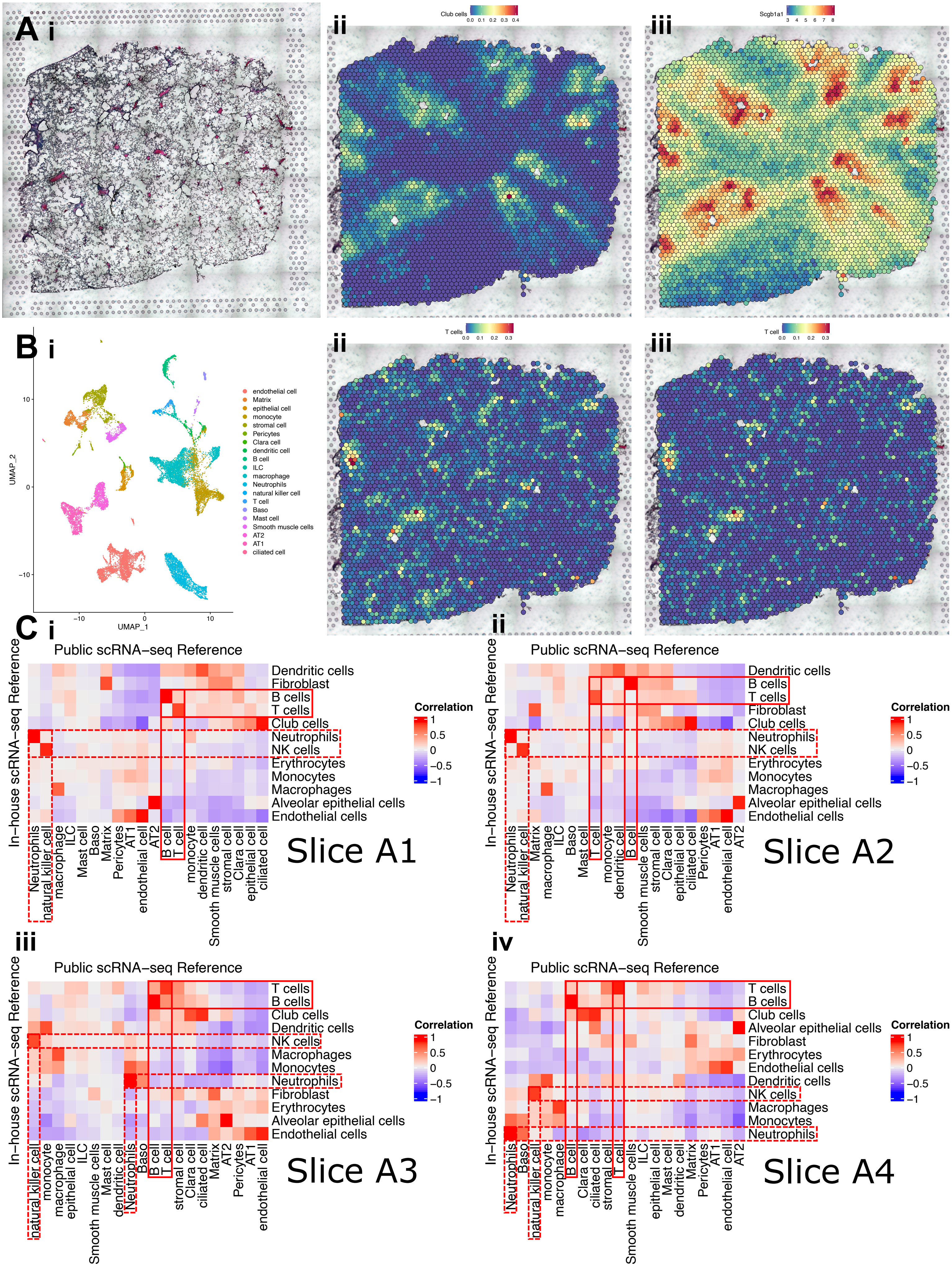

### ST_figure_3.pdf

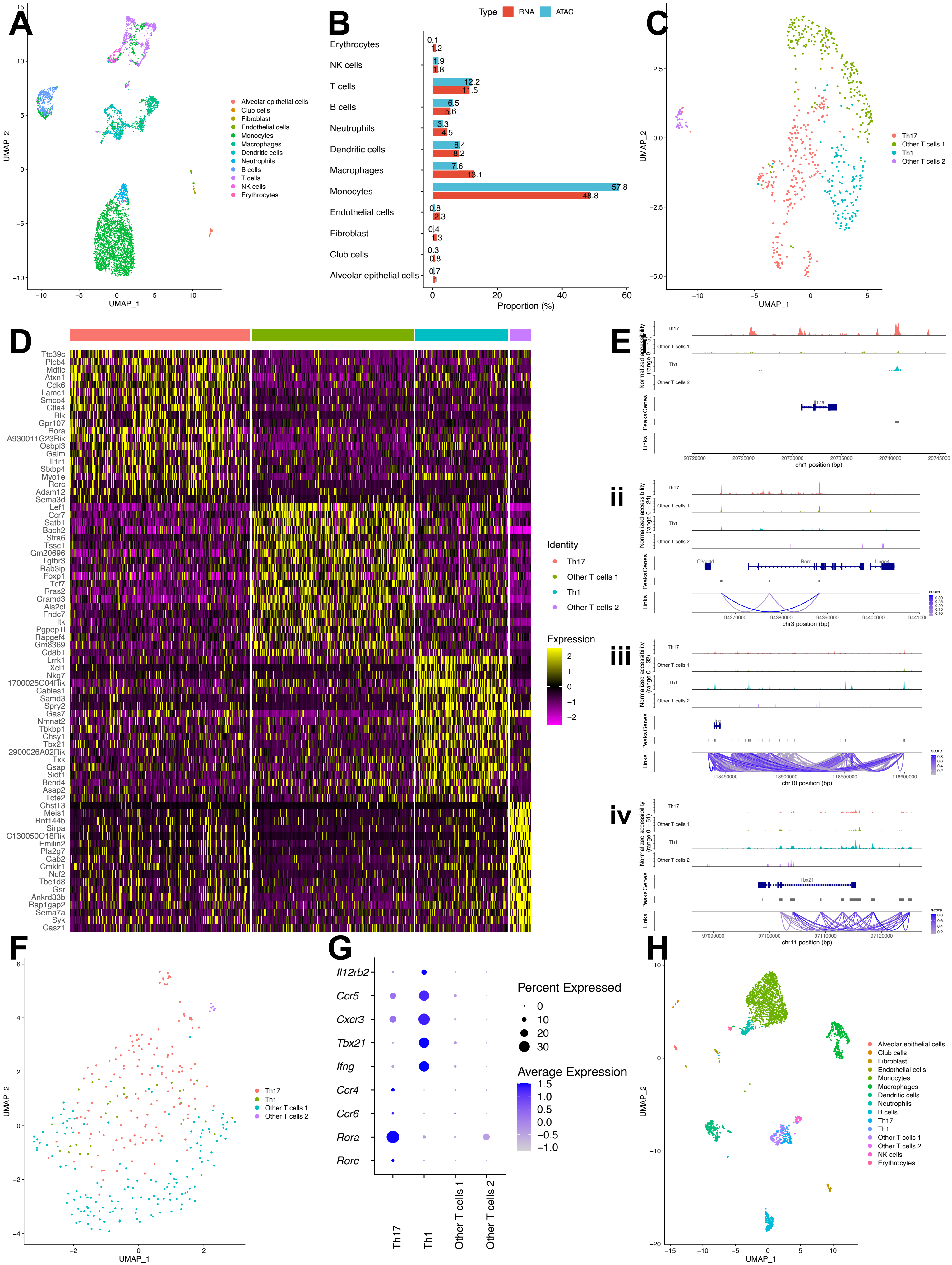

### ST_figure_4.pdf

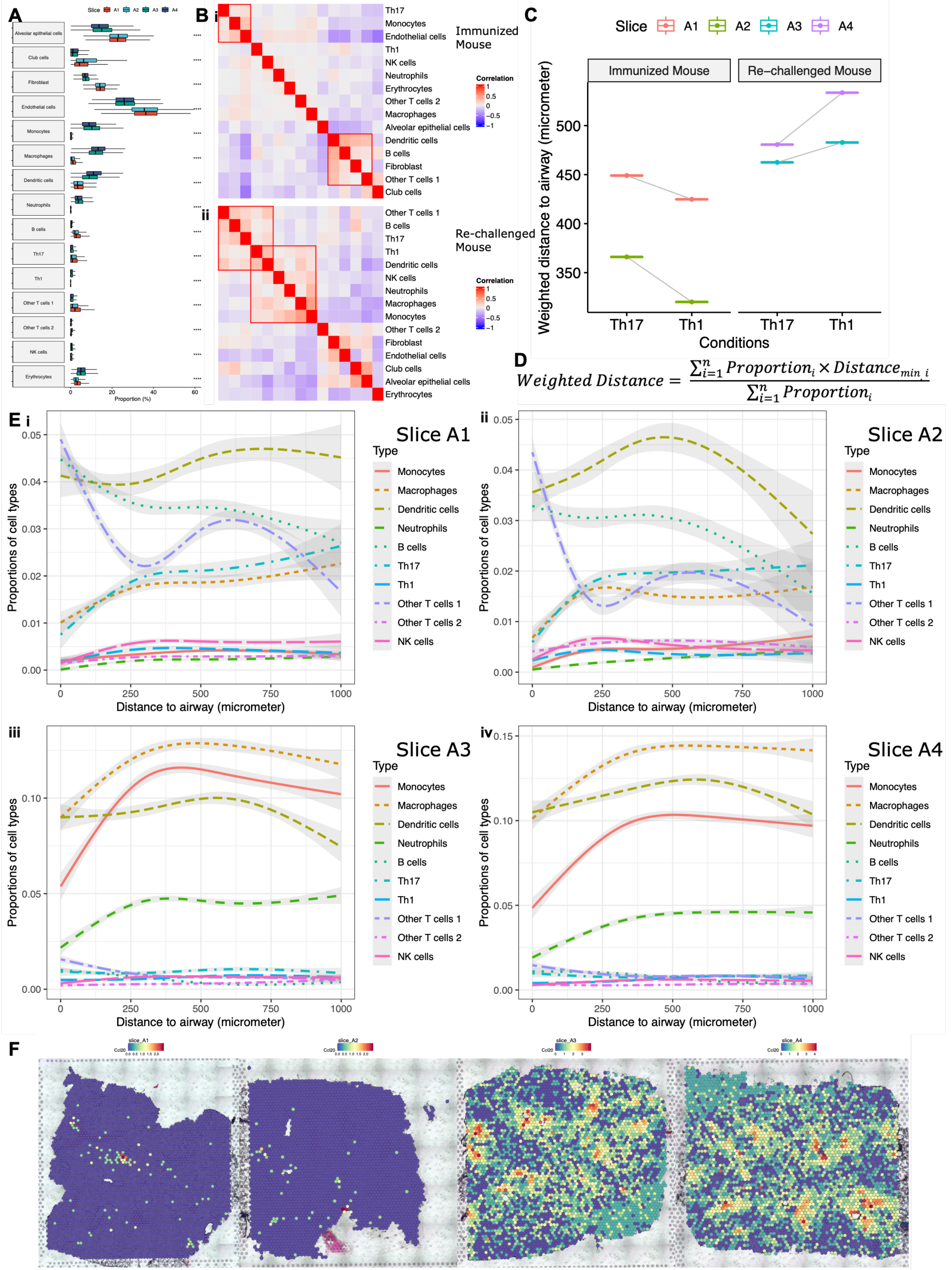

### ST_figure_5.pdf

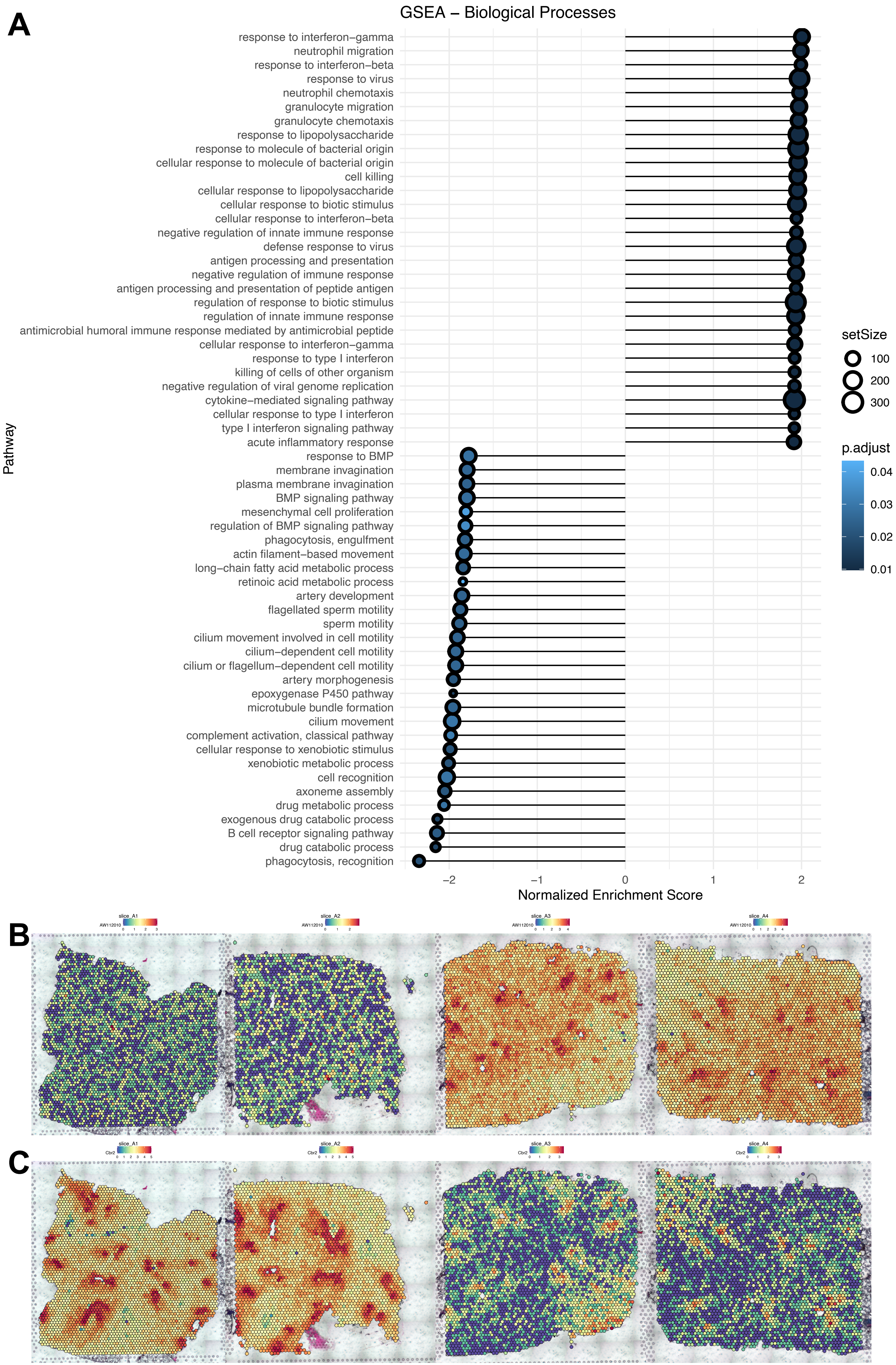

### ST_figure_6.pdf

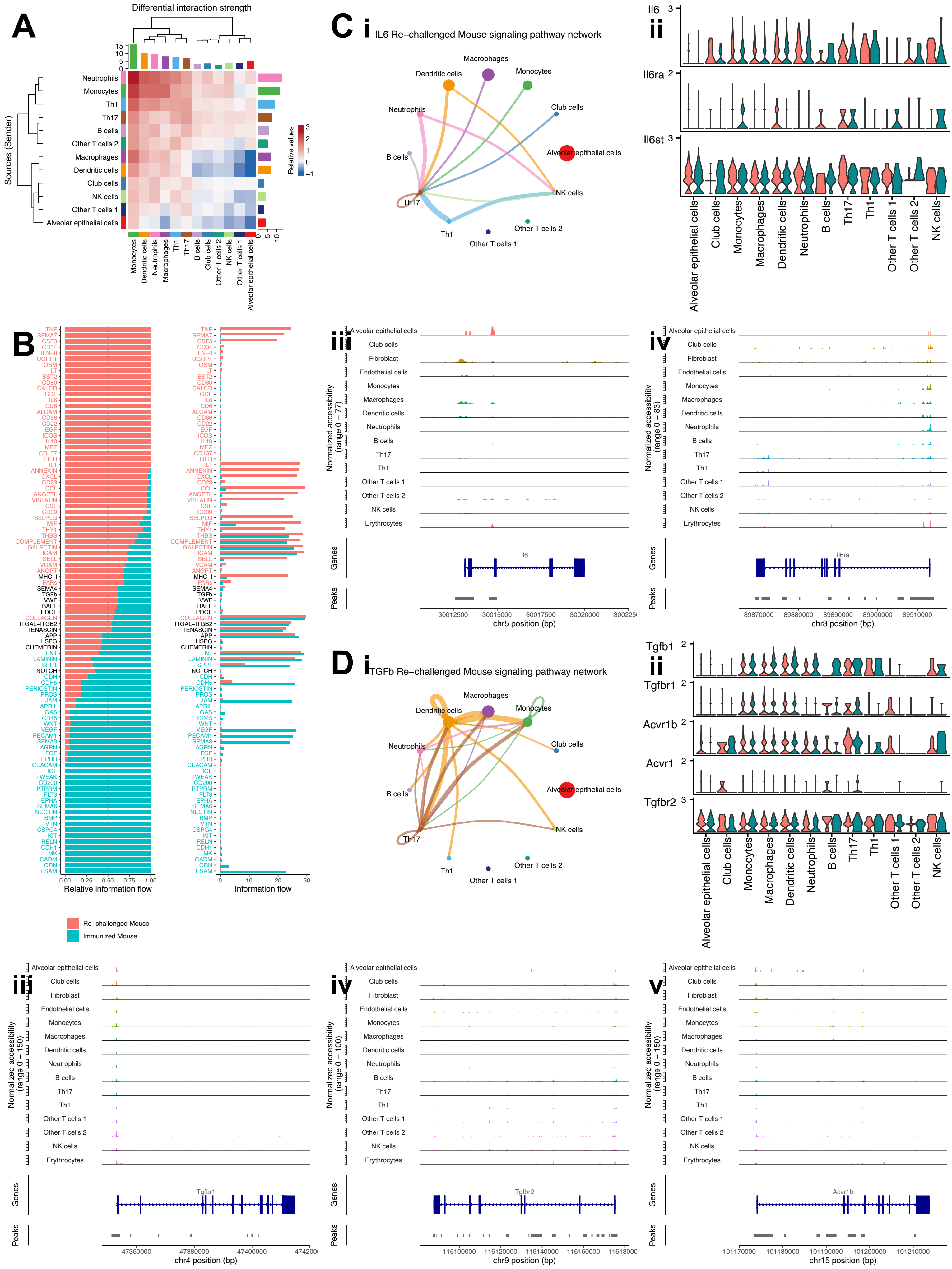

### ST_figure_S1.pdf

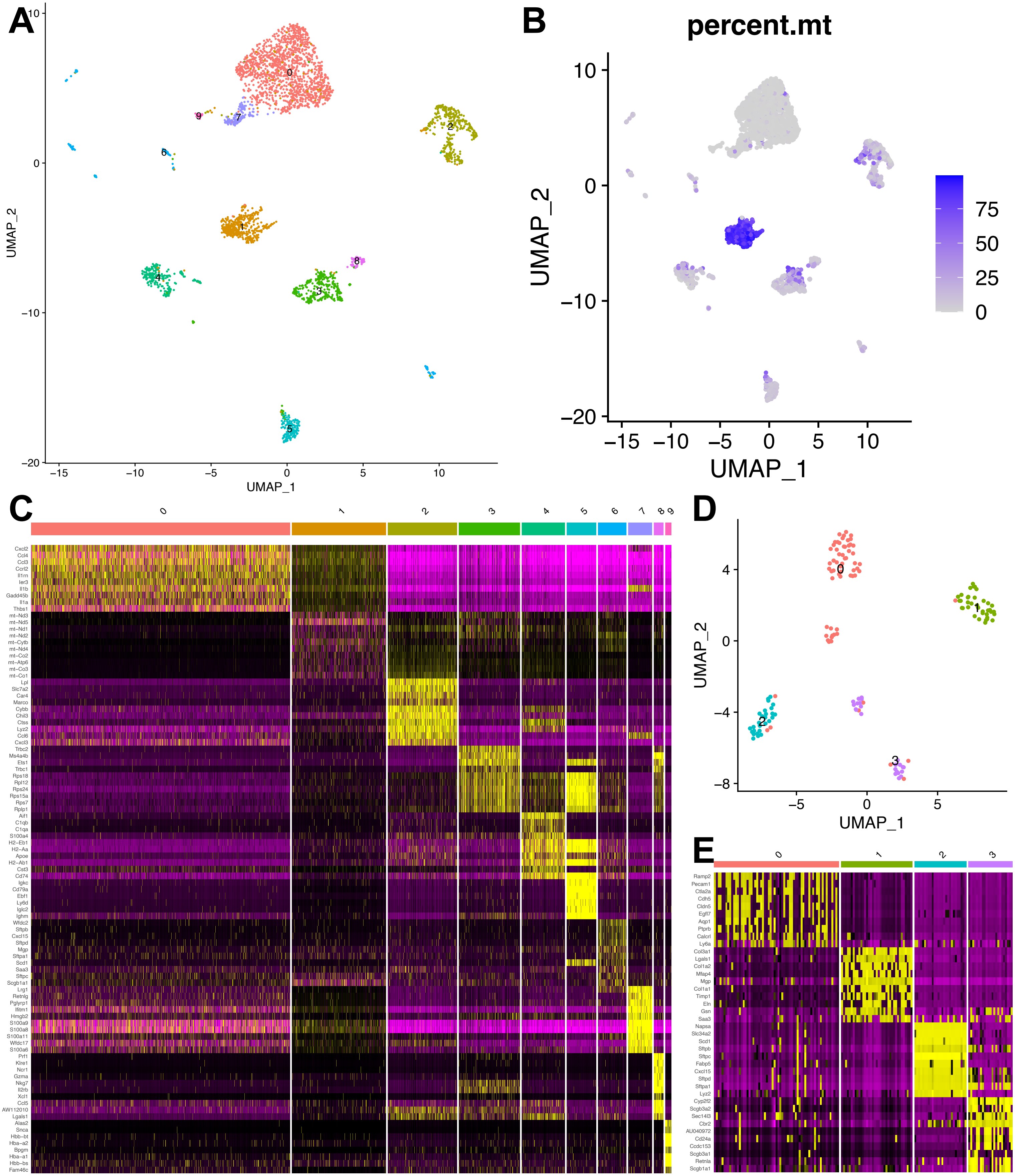

### ST_figure_S2.pdf

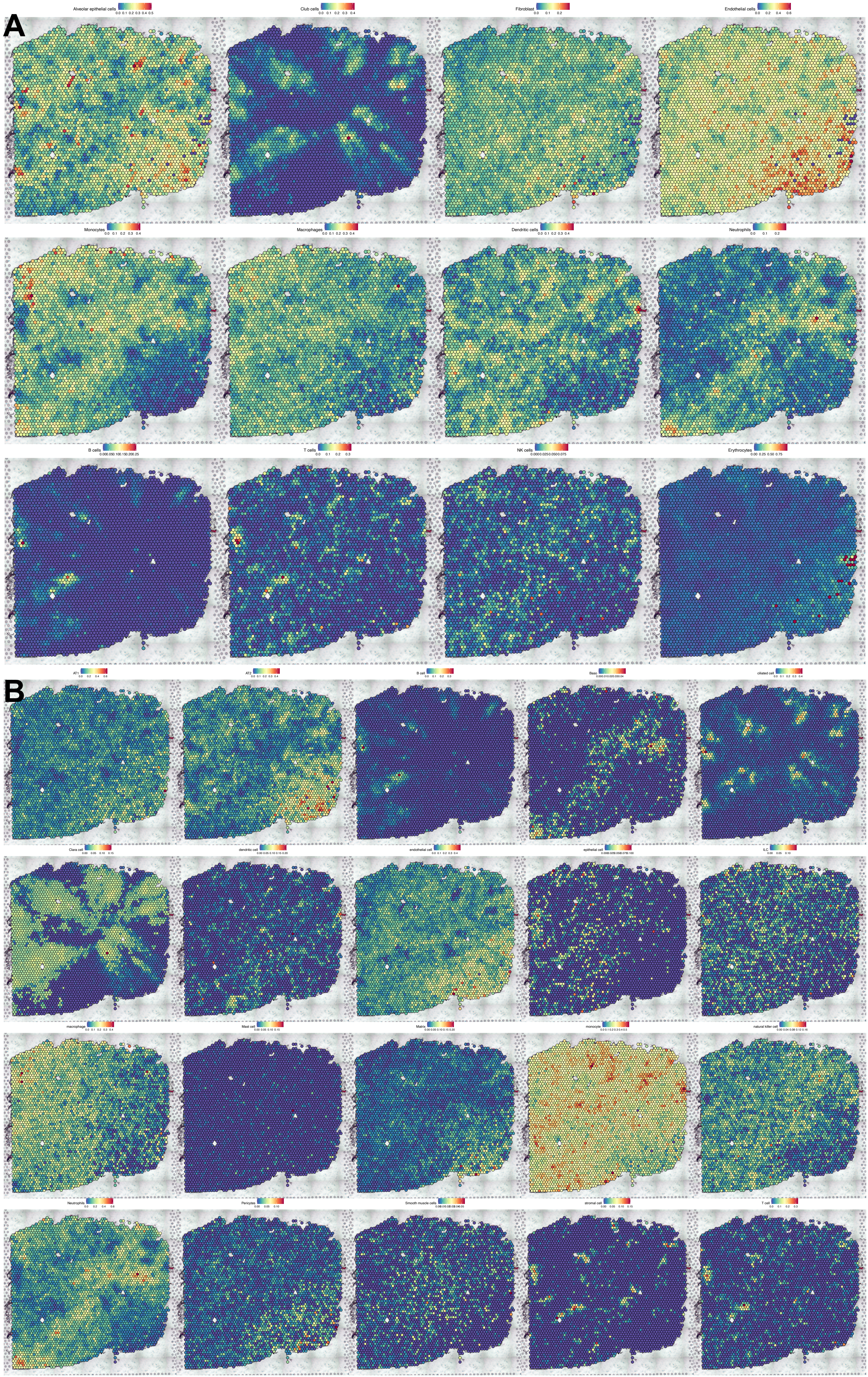

### ST_figure_S3.pdf

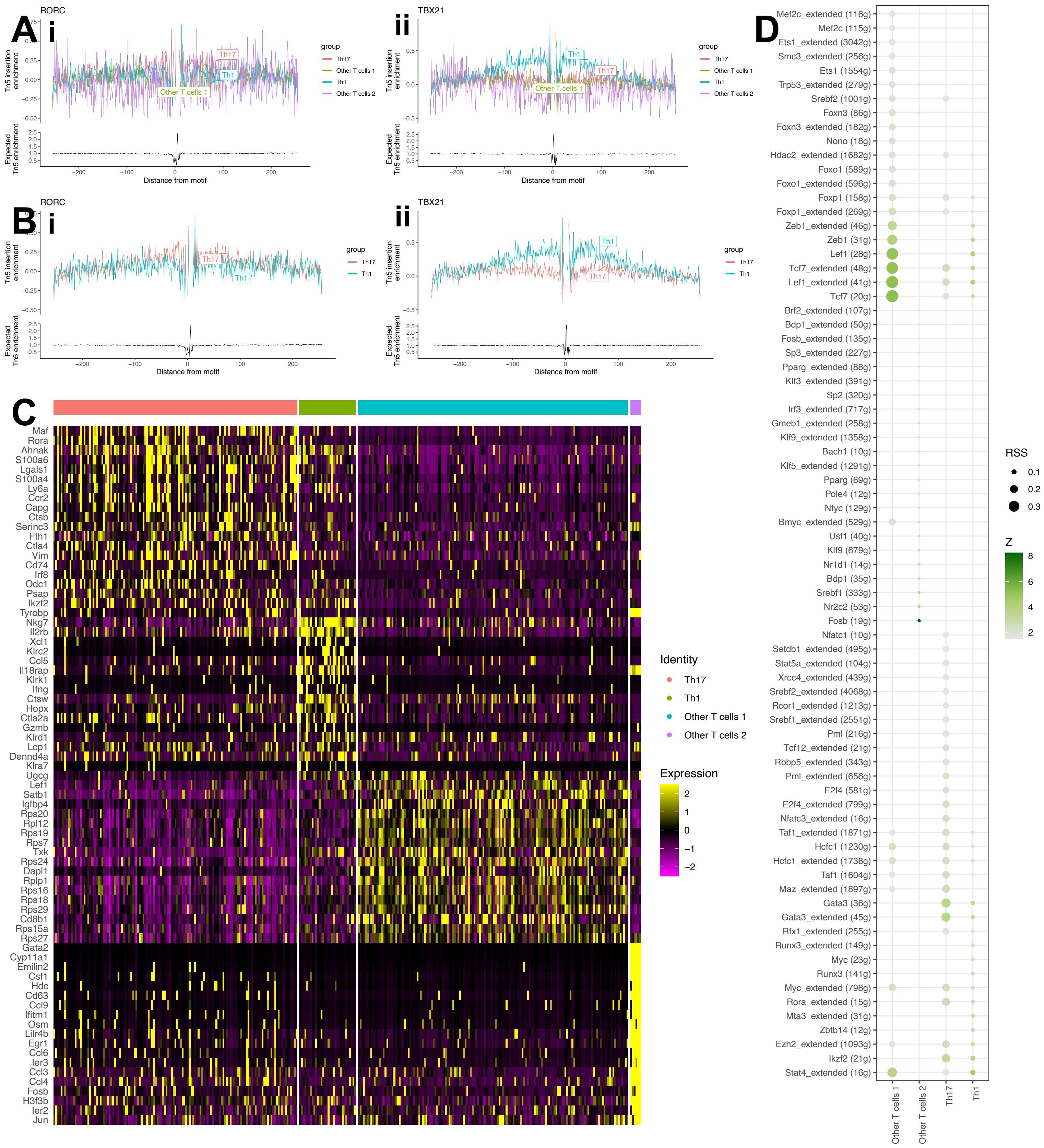

### ST_figure_S4.pdf

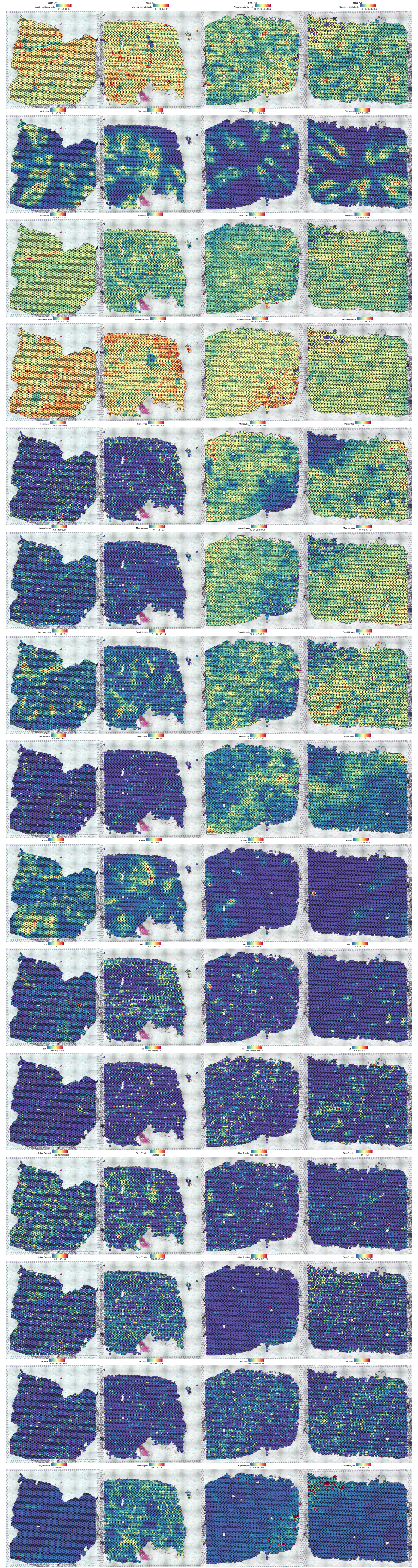

### ST_figure_S5.pdf

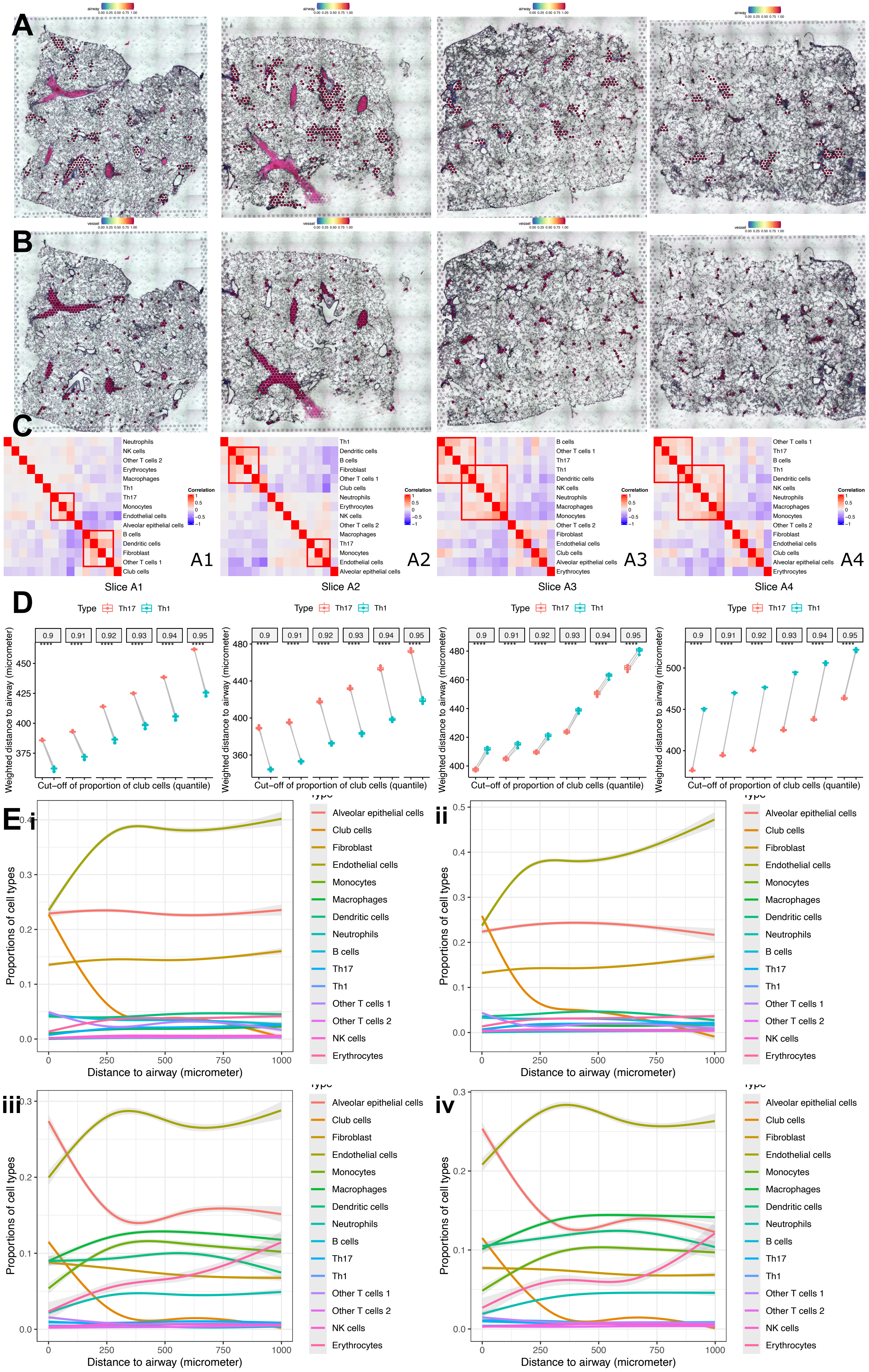

### ST_figure_S6.pdf

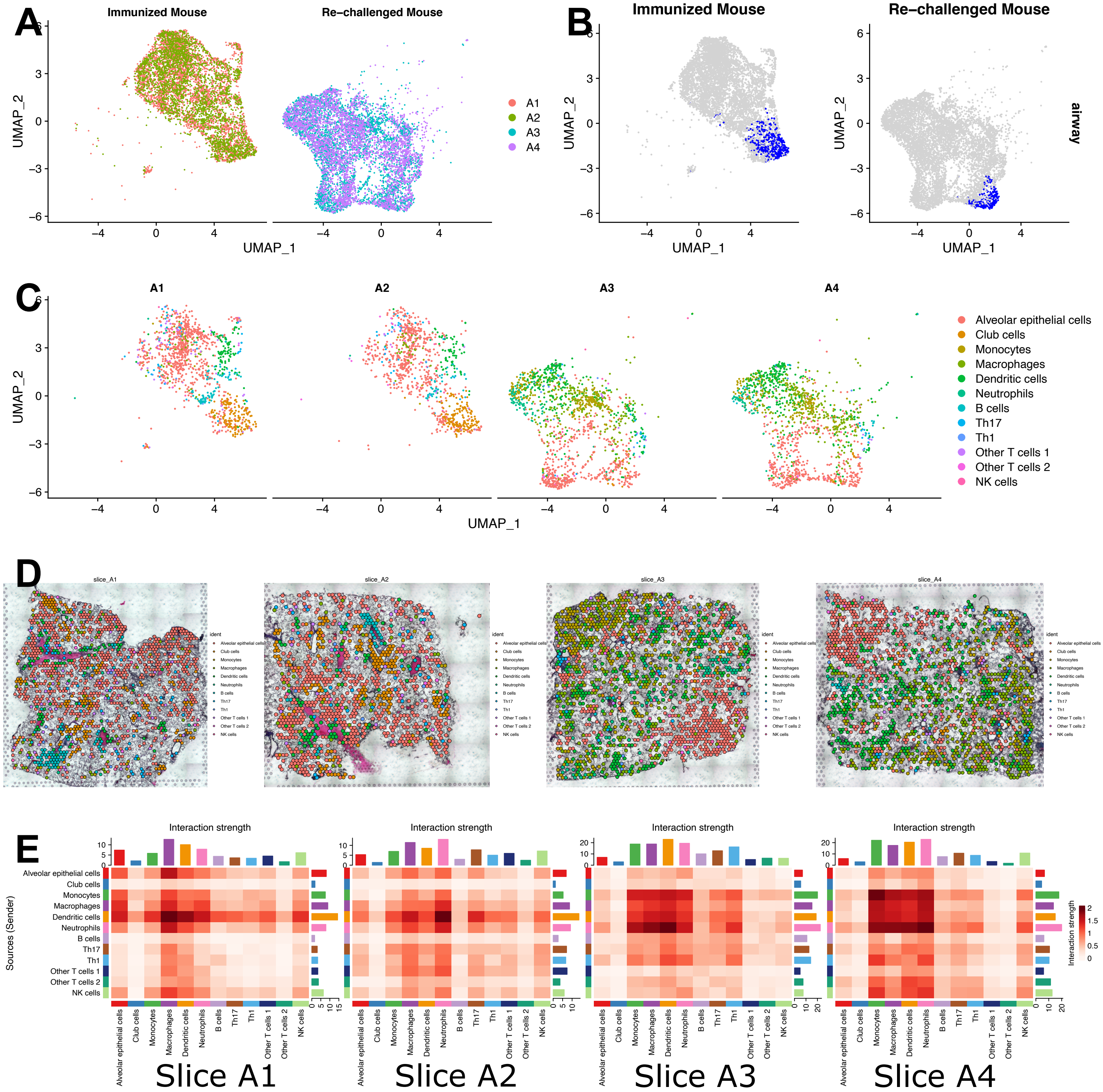
